# A trapped double bond-photoisomerization intermediate in a bacterial photoreceptor

**DOI:** 10.1101/155374

**Authors:** Xiuling Xu, Astrid Höppner, Kai-Hong Zhao, Wolfgang Gärtner

## Abstract

**Abbreviations:**

ASU: asymmetric unit
BV, PCB, PVB (bilin compounds serving as chromophores): biliverdin Ixα, phycocyanobilin, phycoviolobilin
CAPSO: *N*-cyclohexyl-2-hydroxyl-3-aminopropanesulfonic acid
CBCR: cyanobacteriochrome
GAF (protein domain): c**G**MP-specific phosphodiesterases **a**denylyl cyclases and **F**hlA
IMAC: immobilized metal-affinity chromatography
MR: molecular replacement
PAS (protein domain): Per-Arnt-Sim
PHY (protein domain): phytochrome-specific
Pfr, Pg, Pr: far red-, green-, and red-absorbing states of phytochromes and CBCRs

**Summary:** The GAF3 domain of cyanobacteriochrome Slr1393 (*Synechocystis* PCC6803) with an *in vivo* assembled phycocyanobilin (PCB) chromophore has been crystallized in parental state (1.8 Å) and photoproduct state (1.86 Å), identified by 15-*Z* and 15-*E* chromophore configuration. Comparison of both structures for the same protein allows precise determination of structural changes after photo-activation. The chromophore photoisomerization causes an outward movement and partial helix formation of a formerly unstructured loop. A tryptophan residue located in this loop, in π-π stacking distance to PCB in the dark state, moves away by 14 Å opening the binding cleft for the entry of water molecules. Also the *in vitro* assembled protein (chromophore addition to apo-protein) has been crystallized (1.6 Å resolution). Most importantly, an intermediate structure was solved (2.1 Å) with the protein in photoproduct conformation and the chromophore already isomerized into the parental 15-*Z* configuration, thereby giving insight into chromophore-initiated conformational protein changes.

**Impact Statement:** This manuscript presents crystal structures of a photochromic protein in both states, before (1.6 Å) and after (1.9 Å) the light induced photochemical event with sufficient resolution to allow detailed description of conformational changes of chromophore and protein. The light driven reaction, double bond photoisomerization of a covalently bound bilin chromophore is presented here for the first time. Our results allow determining the impact of the chromophore photochemistry on the protein conformation. In addition, we succeeded in trapping an intermediate carrying the chromophore already in isomerized state with the protein still in unchanged conformation. Absorption spectra of this intermediate clearly demonstrate a color change, thus allowing conclusion that the absorption of phytochromes is predominantly determined by the chromophore conformation alone with only moderate effect of the surrounding protein.

**Authors’ Contributions:** XX, KHZ, and WG designed the experiment. XX generated the protein. AH performed crystallization trials, collected the X-ray diffraction data and solved the structure. All authors contributed in preparing the manuscript.

## Introduction

Cyanobacteriochromes (CBCRs) constitute a subgroup of the superfamily of bilin-binding, phytochrome-related photoreceptors (Ikeuchi and Ishizuka, 2008; Rockwell and Lagarias, 2010; Anders and Essen, 2015). The bilin-type chromophores comprise phytochromobilin (in plant phytochromes), phycocyanobilin (PCB, in canonical cyanobacterial and CBCR-type phytochromes), and biliverdin IXα (BV, in bacteriophytochromes). As a common feature, all these proteins present a bilin lyase activity catalyzing formation of a covalent thioether bond between the chromophore and a conserved cysteine residue, and they all show photochromicity, i.e., a shift in absorbance between the parental or resting state and the photoproduct. The best-known representatives of this photoreceptor family are canonical phytochromes, e.g., those from higher plants, being formed *in vivo* as a red-absorbing parental form (Pr) that converts upon irradiation into a far red-absorbing photoproduct (Pfr) (Montgomery and Lagarias, 2002). The photoproduct’s thermal stability of canonical phytochromes, once formed, varies from hours to minutes for individual phytochromes. In these ‘classical’ phytochromes a PAS, a GAF, and a PHY domain arrangement (with only very few exceptions of either PAS or PHY domain missing) is essential to maintain the spectral and kinetic properties. In contrast, however, CBCRs unite all these properties within a single GAF domain, and in fact, the chromophore-bearing GAF domain of CBCRs maintains all these features (lyase activity, photochromicity) even when expressed autonomously as a small protein (Narikawa et al., 2008; Zhang et al., 2010).

Due to these features, CBCRs have attracted scientific interest as potential protein tags and tools in optogenetics applications (Zhang et al., 2010; Simon et al., 2017) considering their small size and their photochemical properties: whereas canonical phytochromes switch between a red- and a far red absorbing state, the absorbance maxima of CBCRs (resting states and photoproducts) cover the entire visible spectrum, even expanding into the near-ultraviolet range (Rockwell et al., 2008; Ikeuchi and Ishizuka, 2008; Rockwell et al., 2012). In addition, CBCR-GAF domains show a noticeable fluorescence for both photoswitchable states ( Zhang et al., 2010; Chen et al., 2012; Ma et al., 2012; Simon et al., 2017) which is virtually absent in canonical phytochromes.

Reversible conversion between parental and photoproduct state in CBCRs - as in canonical phytochromes - is initiated by irradiation of the bilin chromophore with light of appropriate wavelength. The key event in photoconversion is a double bond photoisomerization of the chromophore in which the protein constrains the photochemistry to the double bond between rings C and D of the bilin (Figure 1 a). The photoisomerization proceeds through a conical intersection and is completed within few picoseconds (Müller et al., 2008; Kim et al., 2012b; Kim et al., 2012a; Slavov et al., 2015), followed by protein motions enforced by the changed chromophore conformation that expand into the long milliseconds time range (Fukushima et al., 2011; Xu et al., 2014). The protein conformational changes finally generate a biological signal allowing the organism to adapt to the detected illumination conditions.

**Figure 1:**
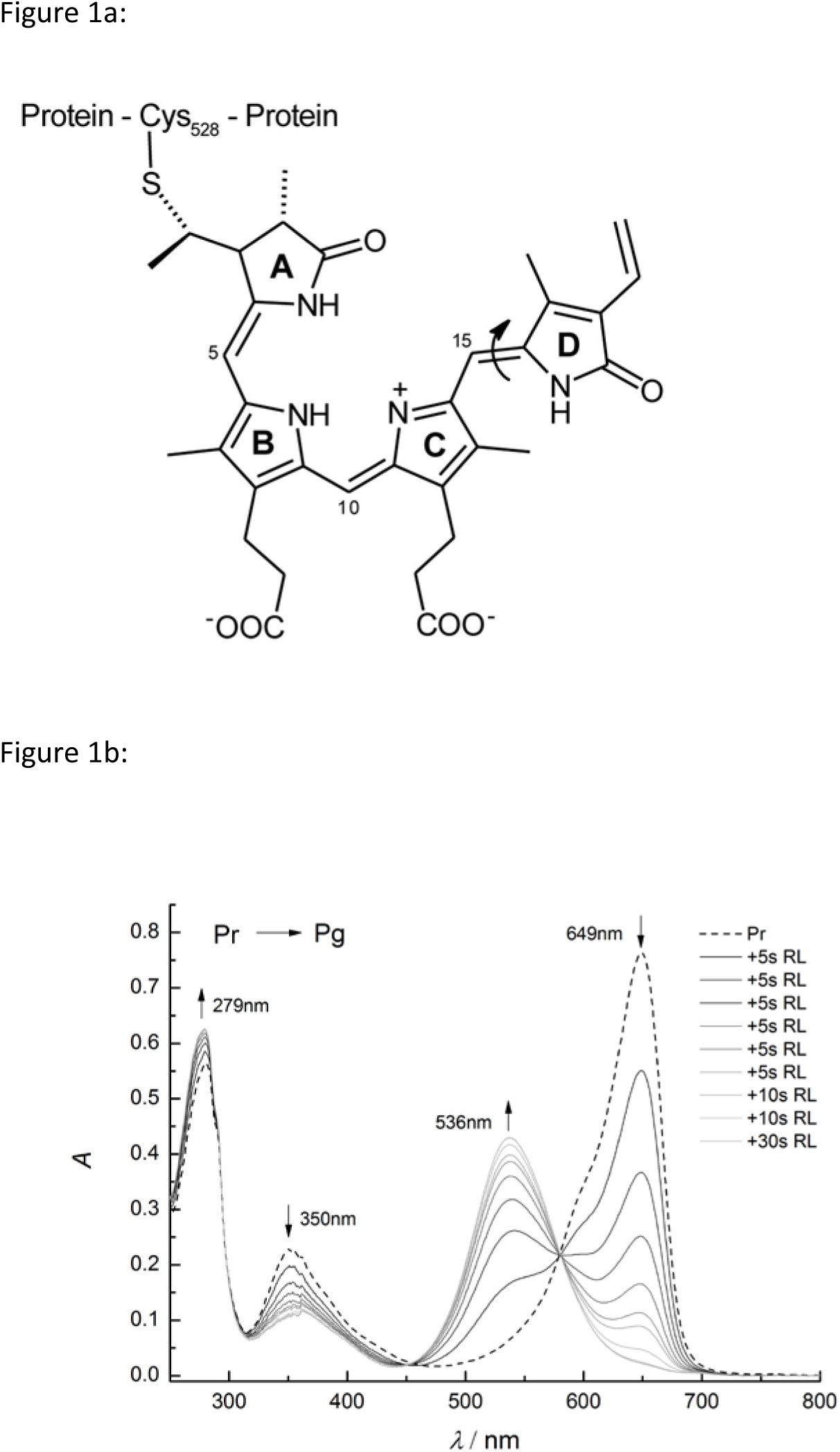
a) Chemical structure of protein-bound phytochromobilin chromophore of Slr1393g3. Attachment to the protein in Slr1393g3 is accomplished through a thioether bond between Cys528 and the 3^1^ position of PCB. The molecule is shown in the parental state configuration (*Z,Z,Z,s,s,a*). Rings A – D are labelled, and the double bond photoisomerization (double bond between rings C and D) is indicated by an arrow. (b) Absorption spectra of both red- and green-absorbing forms of Slr1393g3. Formation of the green-absorbing form is shown upon stepwise irradiation of the red-absorbing form (total irradiation time 80 s, irradiation source 670 nm LED).

Cyanobacteriochromes have been classified into several sub-families according to their absorbance range and amino acid sequences ( Rockwell et al., 2009; Rockwell et al., 2011; Anders and Essen, 2015). The most prominent subfamily is red-green switching with a red-absorbing parental state (absorption maximum around 650 nm) and green-absorbing photoproduct showing a remarkably large hypsochromic shift of *ca*. 100 nm. Slr1393 from *Synechocystis* PCC6803 is a member of this sub-family with absorbance maxima at λ_max_ = 649 nm and 536 nm, respectively (Chen et al., 2012; Xu et al., 2014). Slr1393 is composed of three consecutive, N-terminally located GAF domains and a C-terminally located histidine kinase, making this protein a canonical two component signaling system (Chen et al., 2012). Out of the three GAF domains, only GAF3 (Slr1393g3, in the following,…gn, e.g., g3, indicates the number of the GAF domain of a CBCR that binds the chromophore) has the capability to bind a bilin chromophore, however, this property, as well as the photochemical features, are also preserved if Slr1393g3 is expressed independently as an autonomous protein (Figure 1 b) (Xu et al., 2014). The absorbance maxima of both parental and photoproduct state are sufficiently well separated allowing a nearly complete mutual light-driven conversion of one form into the other, and, in addition, the photoproduct state of Slr1393g3 shows a remarkable thermal stability: in the dark, the thermal re-conversion into the parental state proceeds to less than 10 % over 24 h (20 °C) (Xu et al., 2014).

These properties allowed crystallization and structure determination of both parental state and photoproduct spectral forms of Slr1393g3 with high resolution, enabling us to precisely identify conformational changes between parental state and photoproduct in one and the same protein. So far, nearly all approaches in phytochrome structure analysis aiming at the understanding of conformational changes between the parental and the photoproduct state had to rely on comparison of the two states from different proteins for which one or the other state is the parental form, e.g., comparing a bacteriophytochrome (15-*Z* red-absorbing parental state) with a ‘bathy’ phytochrome for which the 15-*E* far red-absorbing form is the parental state (Yang et al., 2008). Only one other phytochrome-related protein, the CBCR TePixJg from *Thermosynechococcus elongatus*, has already been crystallized in both parental and photoproduct state with good resolution (2.4 and 2.0 Å) (Narikawa et al., 2013a; Burgie et al., 2013), however, this protein shows a remarkably exotic, non-typical photochemistry. In contrast to canonical phytochromes and red-green switching CBCRs, TePixJg shows a very blue-shifted parental state (λ_max_ = 433 nm) and a green-absorbing photoproduct (λ_max_ = 531 nm). The positions of the absorption bands of both states of TePixJg are caused by atypical chromophores: although being formed with PCB as chromophore, the protein converts this chromophore into phycoviolobilin (PVB, conversion of the C4=C5 double bond, Figure 1a, into a single bond), thus generating a chromophore with only three pyrrolic rings (B, C, and D) in conjugation (Ishizuka et al., 2011), but still allowing *Z*-*E* isomerization at the double bond between rings C and D. Even more uncommon for phytochromes is the chromophoric structure in the blue-absorbing parental state: the short absorption band is accomplished by covalent attachment of a cysteine residue to the central C10 position thus restricting the conjugated π-system to only two pyrrolic rings (rings C and D) (Rockwell et al., 2008).

We thus consider the here presented crystal structures of the parental and photoproduct state of Slr1393g3 an optimally suited paradigm to describe structures of canonical phytochromes and – related proteins.

## Results and Discussion

GAF3 domain of Slr1393 was expressed in *E. coli* as an N-terminally His-tagged protein comprising 155 amino acids. For *in vivo* assembly of the chromophore, expression followed a two-plasmid protocol (Chen et al., 2012; Xu et al., 2014). The affinity-purified protein showed the known spectroscopic features (λ_max_ = 649 and 536 nm for parental and photoproduct state), the nearly complete interconversion of both states (>95 %), and the formerly reported thermal stability of the photoproduct allowing crystallization of both photochromic forms independently (see Materials and Methods Section for details). Crystals of *in vivo* Slr1393g3 in its parental (red-) and photoproduct (green-absorbing) form diffracted up to 1.8 and 1.86 Å, respectively. In addition, the apo-form of the protein, subsequently (*in vitro*) assembled with the PCB chromophore, was crystallized in its parental state and structurally solved with 1.6 Å resolution.

As a most interesting result, a photoisomerization intermediate could be trapped during the re-conversion of the photoproduct into the parental state carrying the chromophore already in the parental 15-*Z* state configuration whereas the protein still exposes the photoproduct conformation. For this intermediate, the structure could be determined with a resolution of 2.1 Å. For data statistics and refinement see Table 1.

**Table 1:** Crystallographic Data Collection and Refinement Statistics of Slr1393g3-Pr, Slr1393g3-intermediate and Slr1393g3-Pg

| Data Collection | Slr1393g3-Pr (*in vitro*) | Slr1393g3-Pr (*in vivo*) | Slr1393g3-intermediate | Slr1393g3-Pg |
| --- | --- | --- | --- | --- |
| PDB code | 5DFY | 5DFX | 5M85 | 5M82 |
| Synchrotron, beamline | ESRF, ID23-1 | ESRF, ID29 | DESY, EMBL, P13 | DESY, EMBL, P13 |
| Space group | $P4_12_12$ | $P4_12_12$ | $P6_5$ | $P6_5$ |
| Cell dimensions (Å/°) | a, b = 62.94; c = 119.35; $\alpha$, $\beta$, $\gamma$ = 90 | a, b = 63.12; c = 118.83; $\alpha$, $\beta$, $\gamma$ = 90 | a, b = 75.10; c = 71.21; $\alpha$, $\beta$ = 90, $\gamma$ = 120 | a, b = 75.29; c = 71.38; $\alpha$, $\beta$ = 90, $\gamma$ = 120 |
| Wavelength (Å) | 1.00894 | 0.97916 | 0.9537 | 0.9763 |
| Resolution range (Å) | 44.51 – 1.60 | 44.633 - 1.80 | 65.04 - 2.10 | 65.2 – 1.86 |
| Completeness (%) | 99.8 | 99.68 | 99.6 | 94.4 |
| Observed reflections | 411926 | 152087 | 268681 | 70835 |
| Unique reflections | 32420 | 22947 | 13390 | 18191 |
| Redundancy | 12.7 | 6.6 | 20.1 | 3.9 |
| Wilson *B* factor (Å<sup>2</sup>) | 20.07 | 22.17 | 37.85 | 33.36 |
| $R_{\text{sym}}$ (%) | 0.053 | 0.055 | 0.065 | 0.031 |
| Mean $I/\sigma$ | 25.7 | 19.2 | 33.3 | 25.8 |
| Refinement |  |  |  |  |
| Resolution range (Å) | 33.629 – 1.60 | 19.960 - 1.800 | 65.04 – 2.10 | 65.2 – 1.86 |
| $R_{\text{work}}$ (%) / $R_{\text{free}}$ (%) | 15.46/18.06 | 16.4/19.9 | 17.0/22.2 | 19.3/23.0 |
| Average *B* factor (Å<sup>2</sup>) | 21.81 | 27.0 | 43.35 | 38.39 |
| No. of atoms | 1561 | 1530 | 1457 | 1435 |
| RMSD (bond lengths) | 0.009 | 0.023 | 0.018 | 0.019 |
| RMSD (bond angles) | 1.322 | 2.09 | 1.96 | 1.88 |
| Ramachandran |  |  |  |  |
| Favored (%) | 99.4 | 98.2 | 96.8 | 96.8 |
| Allowed (%) | 0.6 | 1.2 | 3.2 | 3.2 |
| Disallowed (%) | 0 | 0 | 0 | 0 |

### Structure of the red-absorbing parental state of Slr1393g3 (Pr-state)

Structures of the parental state of both the *in vivo* and the *in vitro* assembled GAF3 domain from *Synechocystis* (PDB codes 5DFX and 5DFY) are nearly identical (rmsd 0.13 Å over 158 Cα atoms). Since the resolution of the *in vitro* assembled structure is slightly higher, we refer to this structure in the following. Crystals of *in vitro* assembled Slr1393g3 diffracted to 1.6 Å and contained one monomer per asymmetric unit (ASU). The entire sequence of Slr1393g3 could be modeled into the electron density (residues 441-596 for original full-length numbering of 1393) with two N-terminal residues originating from the expression tag (Gly439 - Ser440). The R-values after refinement were R_work_ 15.5 % and R_free_ 18.1 %, respectively. Slr1393g3 shows a typical GAF domain folding with a central twisted anti-parallel β-sheet sandwiched by α-helices (Figure 2 a). The overall fold is remarkably similar to that from AnPixJg2 (PDB code 3W2Z, (Narikawa et al., 2013b); rmsd 1.87 Å over 154 Cα atoms) (Figure S1 a), and also to a GAF domain structure of a canonical phytochrome (cf. Cph1, PDB code 2VEA, rmsd 1.98 Å over 142 Cα atoms; in the following, names of canonical and bacterial phytochromes, e.g., Cph1, refer to only their chromophore-binding GAF domain) (Figure S1 b) with the individual conformation requirements, i.e. an extended loop structure to adopt the chromophore.

**Figure 2:**
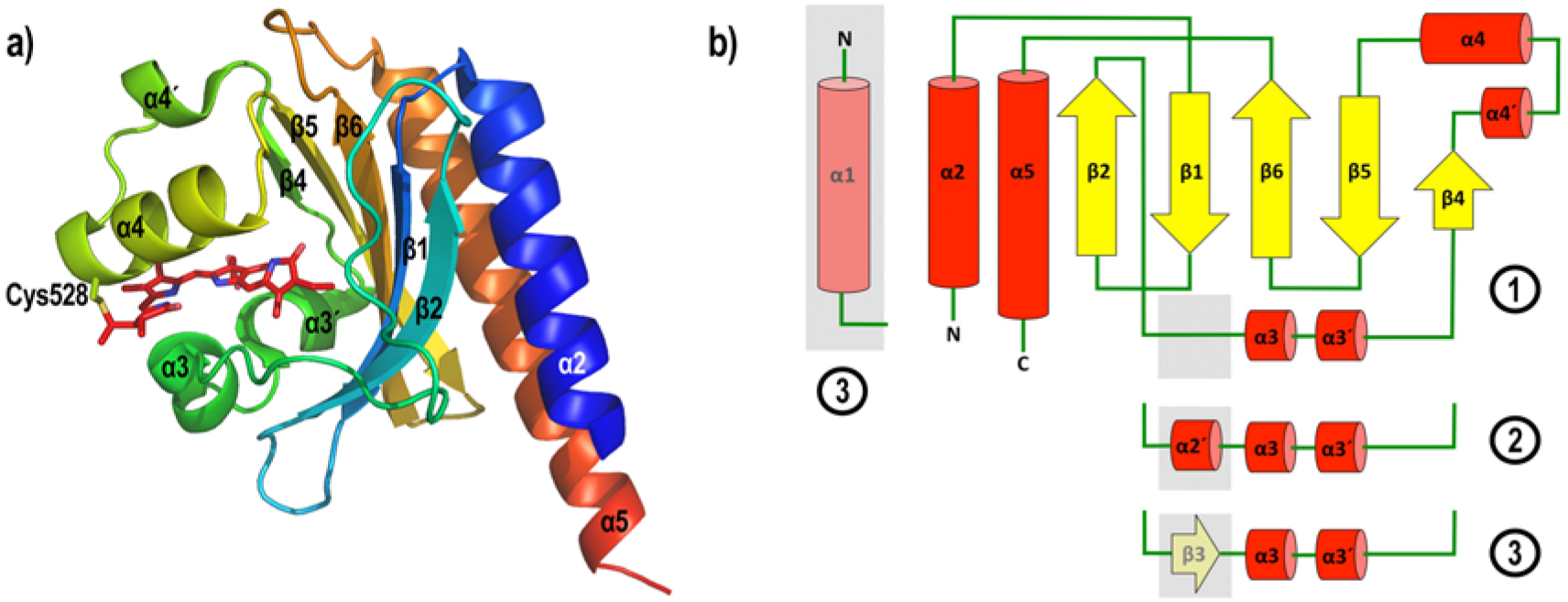
a) Cartoon representation of the overall structure of Slr1393g3 in its *in vitro* assembled parental Pr-state (5DFY). Secondary structure elements are labelled according to the AnPixJg2 structure (3W2Z). The PCB chromophore is shown as sticks, the covalent bond to Cys528 is highlighted. b) Topology of Slr1393g3 in comparison to AnPixJg2: ① Slr1393g3-Pr-state, ② Slr1393g3-Pg and photoisomerization intermediate, and ③ AnPixJg2 topology. Explicitly shown is the Slr1393g3 topology in the parental, red absorbing state. The gray box between β2 and α3 (part of an unstructured loop) converts into a short helical element in the photoproduct and in the photoisomerization intermediate (coined α2’ in the text), and is found as a β-sheet element (β3) in the parental state of AnPixJg2.

The structure of the parental state of AnPixJg2 (3W2Z) and the here reported structure are the only ones known so far for these red-green switching CBCR proteins. In order to allow for facile comparison of secondary structure elements, we refer to the assignment of α-helices and β-sheets as proposed for AnPixJg2 (Narikawa et al., 2013b). Significant variations to the AnPixJg2 structure reside in the absence of β3 (an unstructured loop in Slr1393g3); the here presented additional α-helical element, α4’ (Figure 2 a,b) is also present in the AnPixJg2 structure, but was not annotated by these authors. The central β-sheet is composed of beta strands β1, β2, β4, β5 and β6, flanked by helices α3, α3’, α4’ and α4 on the chromophore binding side, and α2 and α5 on the opposite side.

Leu441 as the first amino acid of Slr1393g3 indicates the beginning of the first helix α2 that extends up to Ser457. A loop (aa 458 – 459) connects this first helical domain with a β-strand (β1, Arg460 – Phe466) being part of an antiparallel arrangement (the second β-strand, β2, covers aa 472 – 479). This first antiparallel sheet region is followed by a large unstructured domain extending up to Gln497 indicating the beginning of two short α-helices α3 and α3’ (Asp498 – Asn504 and Gly506 – His512). A very short β-strand (β4) can be identified for Leu515 – Ala516 – Val517 adapting an antiparallel arrangement with β5 (see below). The following amino acids, Asp519 – Ala523 compose a short helix (α4’). However, except for these very short secondary structure elements, the entire stretch from Ala480 up to Phe525 appears to be a rather unordered loop region. Thr526 – Phe536 form an α-helical motif (α4) harboring the instrumental Cys528 (covalent attachment site for the chromophore). Phe536 as the last aa of this helix is connected through a short unstructured section to Arg539 indicating the beginning of a second antiparallel β-sheet motif (β5: Ala540 – Val548 and β6: Gln551 – Gln560). Amino acids Asn561 up to Trp567 compose a loop connecting this second antiparallel β-sheet motif to the final helix α5 spanning amino acid Gln568 up to Arg594. This final α-helix is oriented antiparallel to the initial helix thus compensating the positively charged N-terminal end by its carboxy terminus.

In Slr1393g3-Pr state the loop connecting β2 and α3 (residues 480 to 497) does not contain secondary structure elements and has two residues that interact with a symmetry related molecule in the crystal assembly (Lys487 with Asn511’ and Gln481 with Glu550’ and Gln551′, respectively) (Figure S2 a). This arrangement is found nearly identical in the AnPixJg2 structure.

The PCB chromophore is covalently bound to Cys528 on helix a4 and is embedded between the central β-sheet array and helices a3, α3’, and a4. As expected for the parental state, the PCB molecule adopt the 15-16 *cis*-configuration, showing an overall *Z,Z,Z,s,s,a* geometry for all double and single bonds in the ring-connecting bridges.

The chromophore is held in its conformation by several interactions to surrounding amino acids (Figure 3, Table 2; note that in the following we use the atom labelling according to the pdb coordinate files when describing inter-atomic distances since this is also the standard labeling in the 2D ligand plots (Figures 3 and S3).

**Figure 3:**
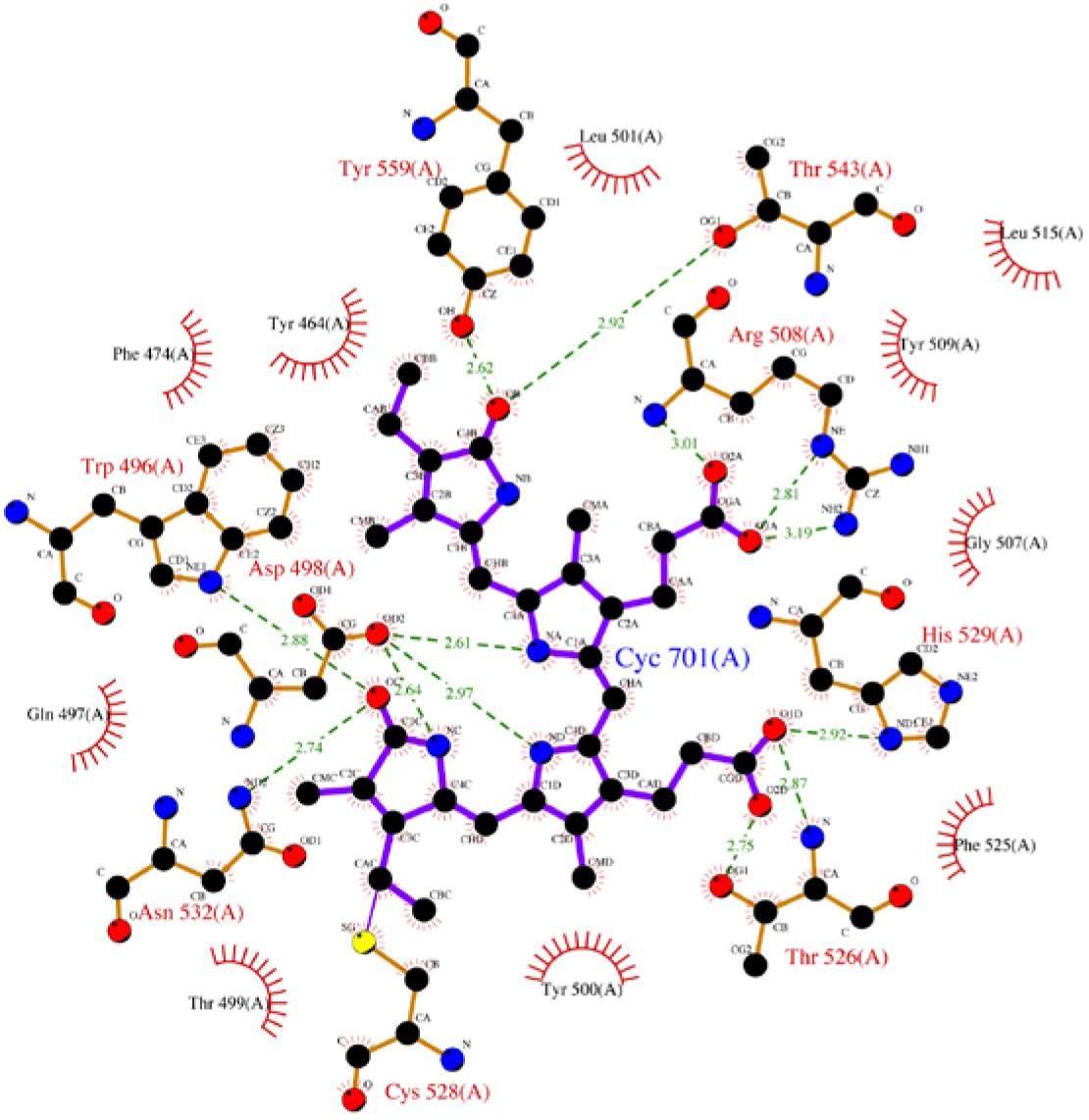
2D ligand plot of the PCB chromophore and surrounding amino acids in Slr1393g3-Pr-state (5DFY). The PCB molecule and interacting residues are represented as ball-and-sticks, distances are given in Å (dotted lines). Residues in close proximity to the chromophore, but not in hydrogen bonding distance are depicted as semicircles, water molecules are omitted. Note that this plot is a purely schematical representation of the interactions and does not reflect absolute chemical configurations.

**Table 2.** Hydrogen bonding pattern of PCB in the various Slr1393g3-states (note that the atom labelling according to the pdb coordinate files are used when describing inter atomic distances since this is also the standard labeling in the 2D ligand plots (Figures 3 and S3), thus enabling easier comprehension).

| PCB<sup>1</sup> | Slr1393g3-Pr *in vitro* assembly (5DFY)<sup>2</sup> | Slr1393g3-intermediate (5M85)<sup>3</sup> | Slr1393g3-Pg (5M82) |
| --- | --- | --- | --- |
| A_OC | Asn532_ND2 (2.74 Å) | Thr499_N (2.93 Å) | Thr499_N (3.05 Å) |
| | Trp496_NE1 (2.88 Å) (π-π stacking with ring D) | - | |
| A_NC | Asp498_OD2 (2.64 Å) | Asp498_OD1 (2.80 Å) | Asp498_OD1 (2.74 Å) |
| B_O1D | Thr526_N (2.87 Å) | Arg508_NH2 (2.69 Å) | Arg508_NE (2.88 Å) |
|  | His529_ND1 (2.92 Å) | CAPSO_NAL (2.59 Å) | HOH (2.68 Å) |
|  |  |  | HOH (2.76 Å) |
| B_O2D | Thr526_N (3.43 Å) | Arg508_NH2 (3.38 Å) | Arg508_NH2 (2.73 Å) |
|  | Thr526_OG1 (2.75 Å) | Arg508_NE (2.90 Å) | CAPSO_NAL (2.69 Å) |
|  | HOH (2.85 Å) | NA2 (2.60 Å) |  |
|  |  | HOH (2.76 Å) |  |
| B_ND | Asp498_OD2 (2.97 Å) | Asp498_OD2 (2.82 Å) | Asp498_OD2 (2.84 Å) |
|  | Asp498_OD1 (3.11 Å) |  |  |
| C_O1A | Arg508_NH2 (3.19 Å) | His529_NE2 (2.84 Å) | HOH (2.64 Å) |
|  | Arg508_NE (2.81 Å) | TRIS_O3 (3.02 Å) |  |
|  | HOH (2.67 Å) | TRIS_O1 (2.39 Å) |  |
| C_O2A | Arg508_N (3.01 Å) | Na2 (2.76 Å) | His529_NE2 (2.73 Å) |
|  | HOH (2.60 Å) |  | HOH (2.71 Å) |
| C_NA | Asp498_OD2 (2.61 Å) | Asp498_OD2 (2.65 Å) | Asp498_OD2 (2.64 Å) |
| D_OB | Tyr559_OH (2.62 Å) | HOH (2.74 Å) | Na2 (2.48 Å) |
|  | Thr543_OG1 (2.92 Å) | HOH (3.22 Å) |  |
| D_NB | HOH (2.68 Å) | HOH (2.94 Å) | - |
|  |  | HOH (3.16 Å) |  |

<sup>1</sup>, first letter refers to the corresponding pyrrole ring system, followed by atom label
<sup>2</sup>, contacts between chromophore and protein are given here only for the *in vitro* assembled protein (better resolution), as both protein structures show virtually the same protein folding and three-dimensional arrangement
<sup>3</sup>, this intermediate originates from the photoproduct state upon X-ray beam exposure and carries the chromophore already in the Z,Z,Z,s,s,a (see text)

The role of an instrumental histidine, considered for chromophore conformational stability and for its absorption properties by π-π stacking (Velazquez Escobar et al., 2013) is taken by His529, located ‘above’ the plane spanned by rings B and C at a distance of 3.8 Å (side chain atom His-ND1 to nitrogen atom ND of PCB ring B) and 3.5 Å (His-NE2 to NA of PCB ring C). Asp498 comprises the central counterion interacting through its carboxylate group (i.e. OD2) with the A-, B-, and C-pyrrolic nitrogen atoms (2.6 Å, 3.0 Å and 2.6 Å to nitrogen NC, ND and NA, respectively). This Asp counterion situation is identically found in AnPixJg2, and is different to, e.g., Cph1 where the central position between pyrrole nitrogens A, B, and C is occupied by the backbone carbonyl group of an aspartate together with a water molecule. In Slr1393g3-Pr state the propionate group at ring B is back-folded making contact to Thr526 (2.8 Å) and to His529 (2.9 Å), whereas the propionate group of ring C is extended and interacts with Arg508 (as well through the side chain as also through the backbone carbonyl unit in a range of 2.8 Å -3.2 Å); the carboxylic group of this propionate makes additional contacts to two closely located water molecules. The carbonyl substituents at rings A and D (OC and OB) are in hydrogen-bonding distance to Asn532 and Trp496 (A-ring carbonyl group), and Thr543 and Tyr559 (D-ring carbonyl group). In addition, a water molecule is part of a hydrogen-bonding network between His529 and D-ring nitrogen (see Table 2; for a comparison of chromophore-protein interactions for the red- (Pr) and the green-absorbing state (Pg), and the intermediate see Figure S3 a-c; for AnPixJg2, Cph1, and TePixJg see Figure S3 d-f).

In particular Trp496 constitutes a stabilizing factor for the Pr-state of Slr1393g3. Its side chain is nearly parallel and in close, π-π stacking distance to ring D (Figure 4) with distances of 3.49 Å between side chain atom CH2 and chromophore atom NB (nitrogen of ring D), and 3.87 Å between CE3 and C2B (carbon 17 in ring D), respectively. This residue undergoes the largest conformational change upon 15-*Z* to 15-*E* conversion (see below). The conformation of the PCB chromophore is clearly identified as *Z,Z,Z,s,s,a* with a nearly planar arrangement of the central rings B and C (-1°), yet with strongly distorted dihedral angles for the A-B and C-D ring arrangements: C4=C5-C6-NB: +24° (corresponding pdb labels: C4C=CHD-CID-ND), NC=C14-C15=C16: -150° (corresponding pdb labels: NA=C4A-CHB=C1B); positive or negative signs indicate clockwise/counterclockwise rotation with respect to the plane formed by rings B and C. By this, rings A and D are tilted upwards the plane determined by rings B and C. The D-ring is kept in place mainly by aromatic amino acids (Tyr464, 3.7 Å; Phe474, 3.6 Å; Trp496, 3.4 Å; Thr543, 2.9 Å; Tyr559, 2.6 Å), and forms an additional interaction through a hydrogen bond to a water molecule.

**Figure 4:**
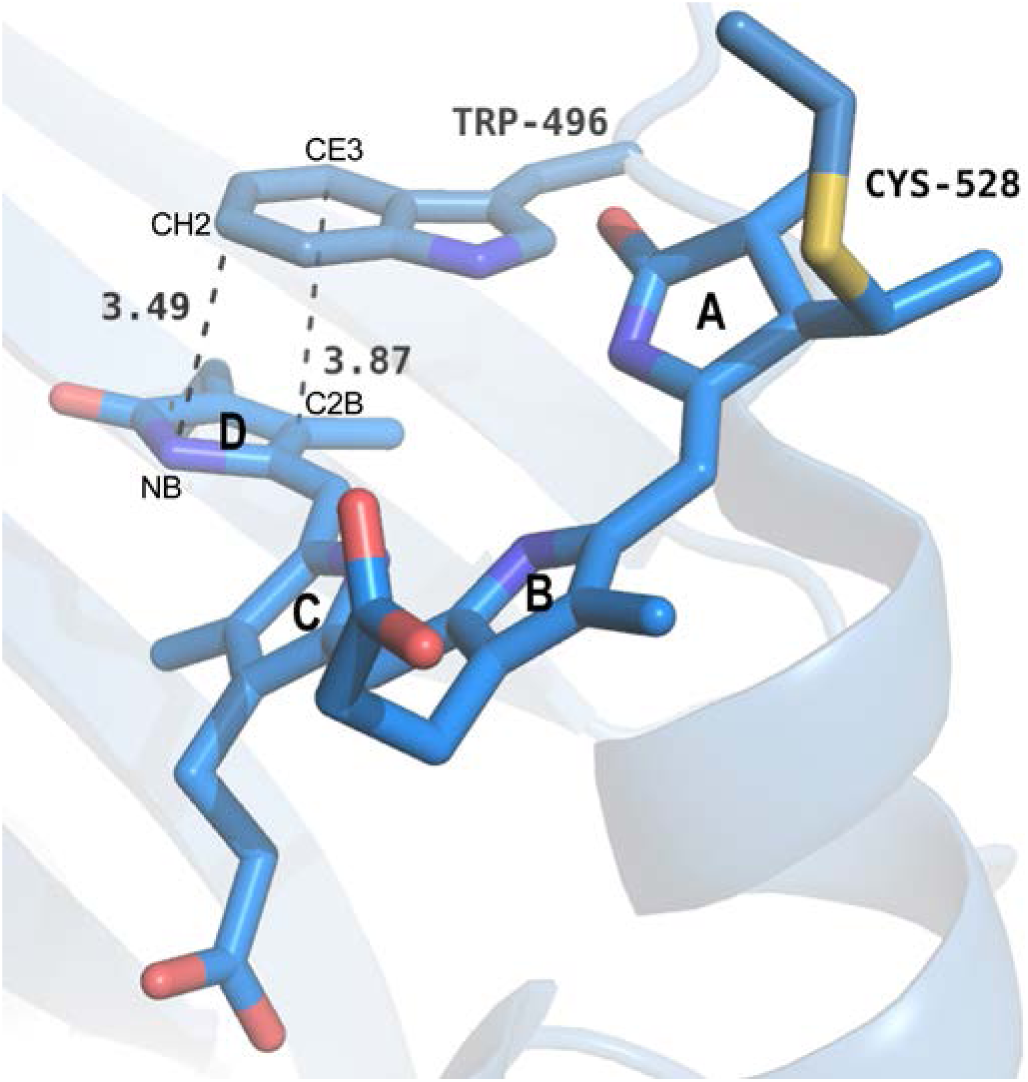
π-π-stacking of Trp496 and D ring of PCB in Slr1393g3-Pr-state. Trp496, the PCB molecule and its covalent bond of to Cys528 are depicted as sticks (helix α4 is omitted for clarity). Distances between atoms (labeled according to the pdb coordinate files, see text) are given in Å.

The chromophore binding pocket is formed as a cleft opening to the protein surface thus enabling the PCB molecule to enter and to be anchored by the covalent thioether bond formation to Cys528. The protein surface charge around the chromophore binding pocket shows an overall acidic character and might be considered as potentially attracting the basic pyrrole rings of PCB prior to covalent bond formation (Figure S4).

### Structure of the green-absorbing photoproduct state of Slr1393g3 (Pg-state)

The thermal stability of the photoproduct allowed crystallization and structure determination of that state with high resolution. Crystals of the photoproduct state of *in vivo* assembled Slr1393g3 diffracted to 1.86 Å and contain one monomer per asymmetric unit (ASU). The entire sequence of Slr1393g3 could be modeled into the electron density (residues 441-596) with one N-terminal (Ser440) and one C-terminal residue of the expression tags (Glu597). The R-values after refinement were R_work_ 19.3 % and R_free_ 23.0 %, respectively. The good quality of the electron density clearly reveals the PCB chromophore being in the *trans*-conformation, i.e. a *Z,Z,E,s,s,a* configuration.

In comparison to the Pr-state of Slr1393g3, most of the secondary structures and the overall protein folding remain the same in the photoproduct structure (rmsd 2.43 Å over 155 Cα atoms). Significant differences are detected in the loop connecting β2 and α3: the connector of β2 and α3 is shifted by *ca*. 8.6 Å towards helix α4 (Figure 5) and part of this loop is re-arranged into a short (two turn) additional α-helix (α2′, Leu486 – Asn492).

**Figure 5:**
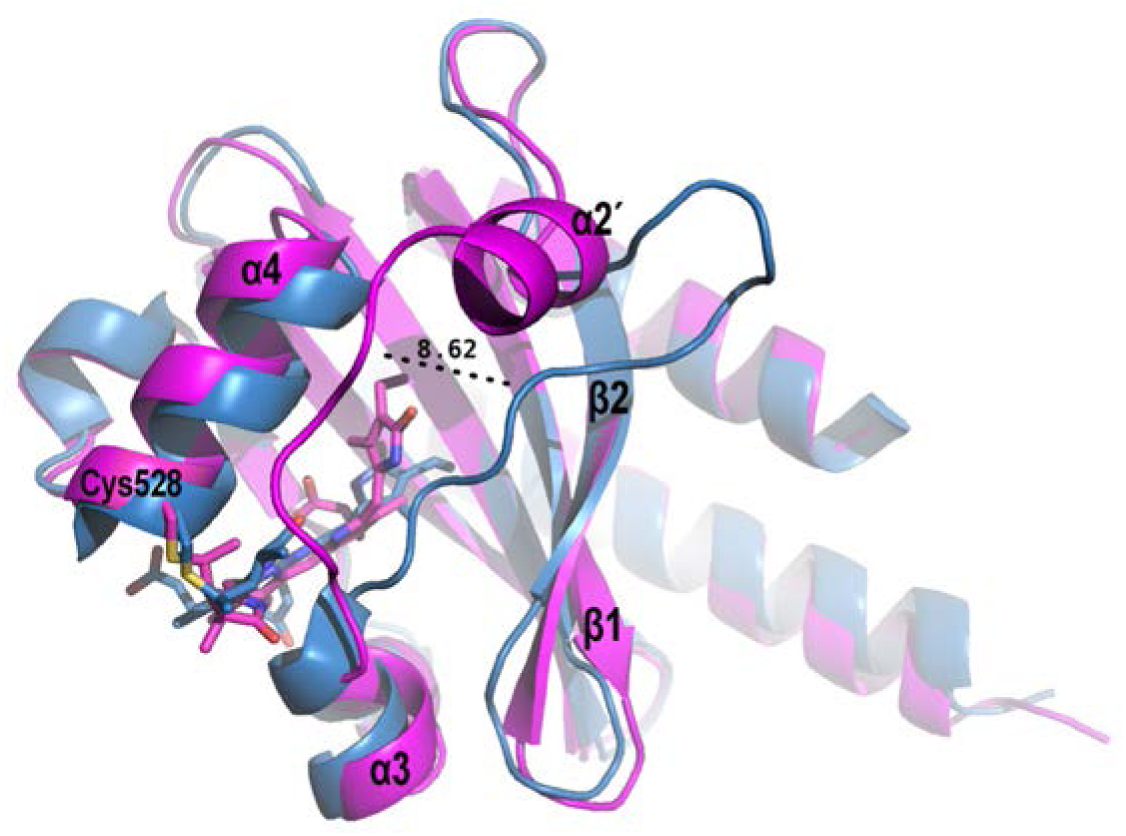
Superposition of Slr1393g3-Pr-state (5DFY, blue) and the photoproduct (Pg) state (5M82, magenta), highlighting the shift of the loop connecting β2 and α2 towards helix α4 and the formation of the additional helix α2’ in the Pg-state.

The formation of this new α-helical element (486-492) adding several hydrogen bonds to this part of the protein might increase stability to the conformation of protein and chromophore in the photoproduct. Some residues of this connecting part are in hydrogen bonding distance to a symmetry related protein molecule (e.g. Gln497 to Glu513′, Leu495 to Arg508’, Asn492 to Gly518’and Arg565′) (Figure S2 b), however, these rather weak interactions are unlikely to be related to the tremendous shift in absorption.

The most significant change in the rearrangement of the long unstructured stretch is an outward swing/rotation of Trp496 (being in π-π-stacking distance to ring D of the PCB molecule in the parental state) by -42° and a displacement of ca. 14 Å with regard to the side chain atom CZ2 (see Figure 6 a). Clearly, the chromophore isomerization had pushed outward this tryptophan residue due to the movement of the pyrrole ring D, thus opening the chromophore binding pocket to the external medium (Figures 5, 6a). As a consequence of the ring D rotation, the chromophore is drawn closer to helix a4 (Figure 6 b).

**Figure 6:**
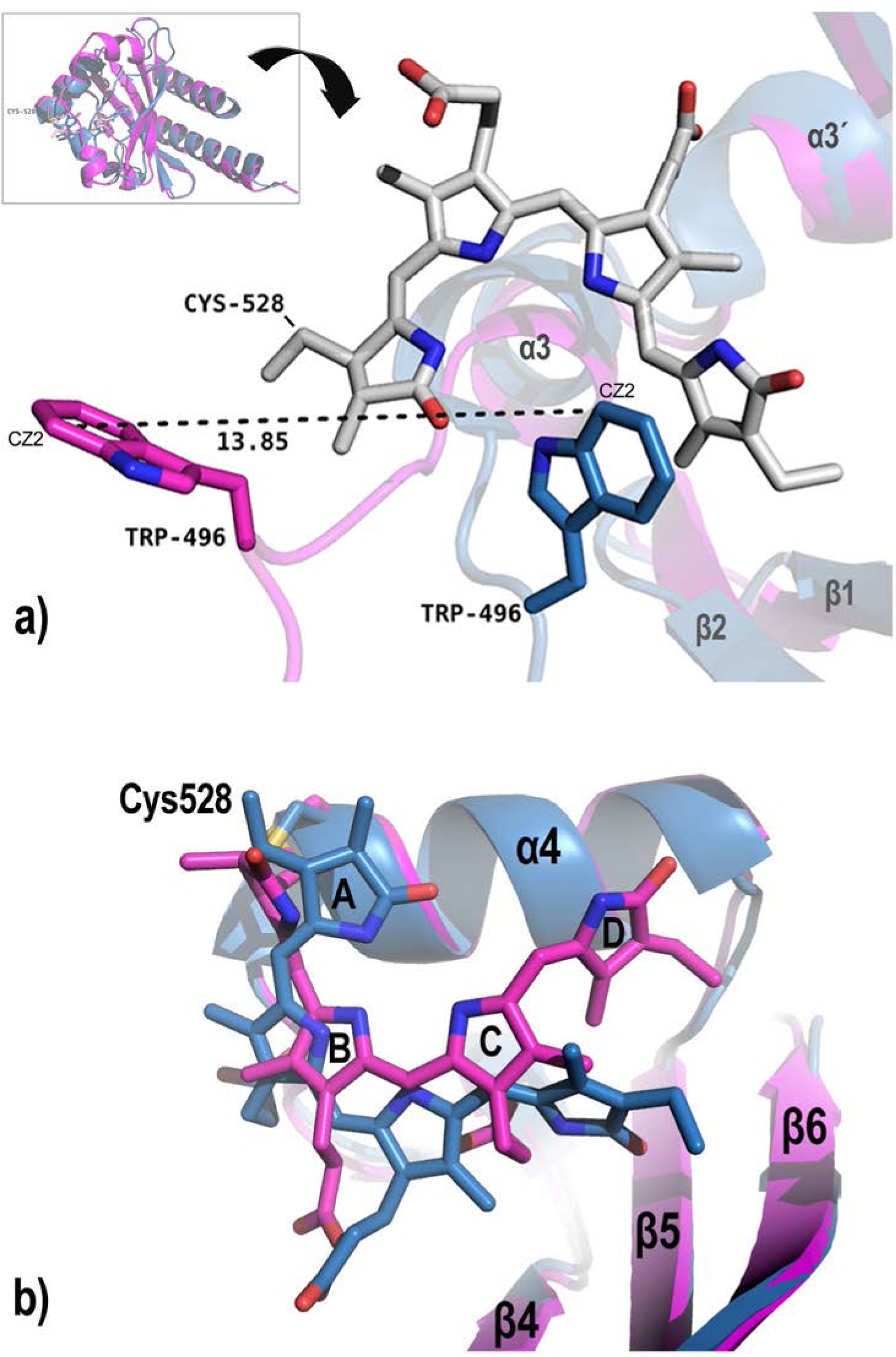
Detailed view of the superposed structures of Slr1393g3-Pr (blue cartoon, 5DFY) and Slr1393g3-Pg (magenta, 5M82). a) Zoom, highlighting the loop and Trp496 movement; the chromophore (gray sticks) is shown in the *Z,Z,Z,s,s,a* configuration of the Pr-state (for clarity the Pg-PCB is omitted), the Pr-Trp is shown in blue, the Pg-Trp is given in magenta. b) Detailed view on the chromophore conformation of both states; the PCB molecule in the Pg-state is, compared with the Pr-state, positioned closer to helix α4 and has moved further into the binding side; color coding as in a).

Further, we find for the Pg-state of Slr1393g3 a change of the propionate side chains’ counterion caused by the chromophore isomerization: still Arg508 keeps this function (~2.8 Å), however, now hand-shaking with the propionate side chain of ring B (photoproduct state) instead of the propionate group from ring C (parental state) (Table 2). The released propionate group from ring C now interacts with His529 (2.7 Å). This change of interaction between Arg508 and either of the two propionate side chains is related to a slight rotation of chromophore around its vertical axis.

As a result of this spatial/rotational re-arrangement, Thr526, Asn532, Thr543 and Tyr559 are no longer involved in chromophore interactions, instead Thr499 stabilizes the A-ring (3.1 Å), together with Asp498 (2.74 Å), as also observed in Slr1393g3-Pr-state (Figure S2 b). Interestingly, a CAPSO molecule, originating from the precipitant, is bound between two monomers close to the propionate group of ring B (see also Table 2), interacting with Gln497 of one chain and Arg508 of the other chain, and therefore further stabilizing the network. In this photoproduct state, four water molecules (2.64 Å, 2.68 Å, 2.71 Å and 2.76 Å) are in hydrogen bonding distance to the oxygen atoms of the propionate groups of ring B and C, whereas only three are present in the Pr-state. Further interaction changes can be seen for ring D: its carbonyl group has contact to sodium 2 (Na2, 2.48 Å), and formerly found (parental state) water molecule is now removed.

The 15-16 double bond isomerization of the PCB chromophore causes distortions along the single bonds between the four pyrrole rings larger than determined for the parental state: The A-ring is much stronger twisted away from the B-C plane (C4=C5-C6-NB: -77°; corresponding pdb labels: C4C=CHD-CID-ND), and even the central bridge between rings B and C experiences a distortion of +8°. The 15-16 double bond isomerization causes again a distortion of the neighboring 14-15 single bond by -120° (NC=C14-C15=C16: -120°; corresponding pdb labels: NA=C4A-CHB=C1B).

### Structure of a trapped double bond isomerization intermediate of Slr1393g3 (intermediate-state)

Extended exposure of above described photoproduct crystals to the X-ray beam yielded an interesting effect such that the protein’s chromophore, crystallized under red light, partly converted from its 15-*E* state into the 15-*Z* state of the parental form, as became obvious from a local color change of the crystal from pink to dark blue at the position of beam exposure. The surrounding protein, however, remained in the photoproduct conformation (see rmsd below) with the features indicative for the photoproduct as described in the former paragraph. A comparison between the chromophore configurations in all three structurally solved states – Slr1393g3-Pr, Slr1393g3-intermediate, and Slr1393g3-Pg – is presented on the basis of their electron density in Figure 7 a,b,c that also highlights the different relative orientation of the chromophore to the surrounding secondary structure elements, e.g., α4 and β5. This color/isomeric change is not of generic nature as would occur from unintended exposure of the crystal to light, but is clearly limited to the location of the X-ray beam, as is apparent by recording the absorbance of the crystal at the position of X-ray beam exposure before and after the data collection conditions (Figure 8): X-ray exposure of Slr1393g3 photoproduct crystals showed an slight increase of the parental form absorbance around 650 nm (absorbance shoulder in Figure 8) with the dominant absorbance still documenting the photoproduct form with λ_max_ around 550 nm. Following that observation, the structure of the photoproduct state (described above) has been determined by very short exposure to the X-ray beam and helical data collection in order to prevent conversion of the chromophore.

**Figure 7:**
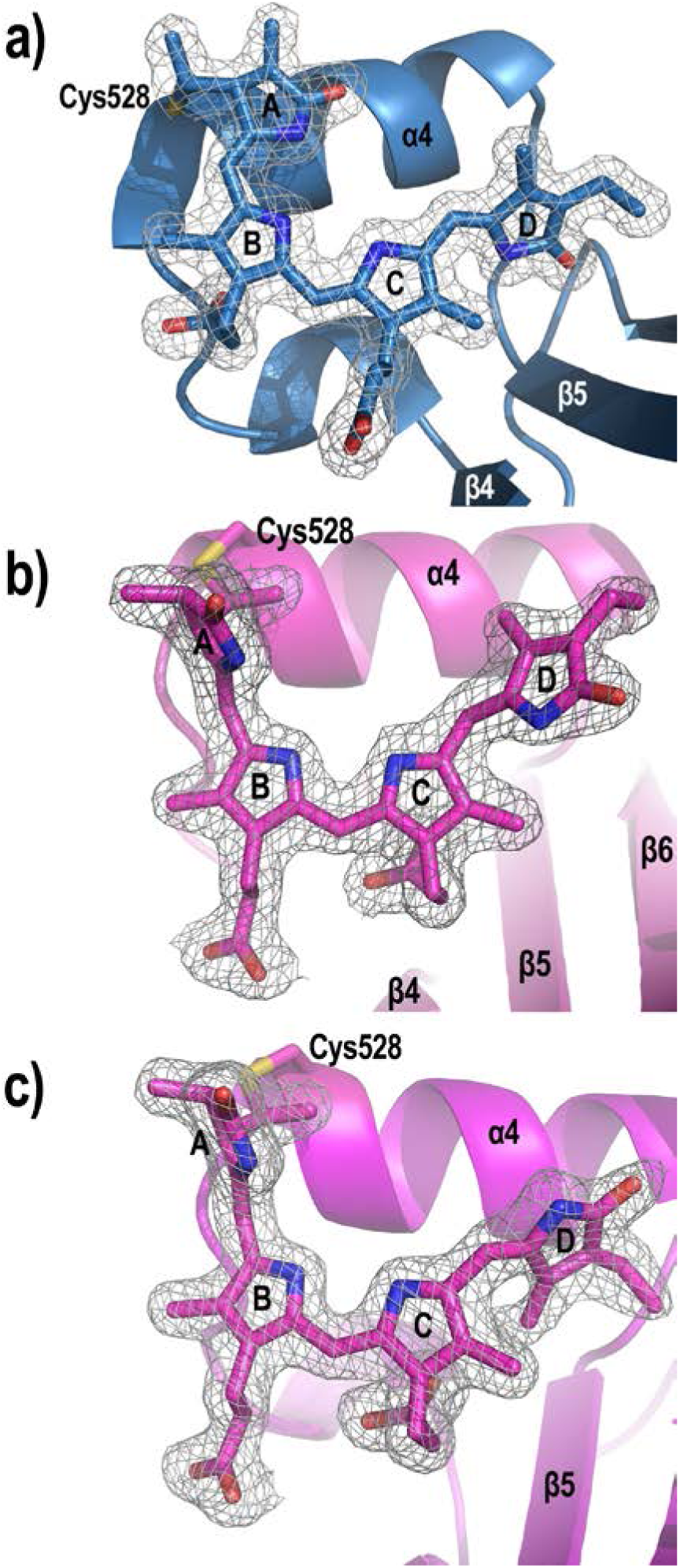
Chromophore conformations of Slr1393g3 in a) the red-absorbing parental state, b) the intermediate (X-ray beam-induced back isomerization of the chromophore) showing a highly distorted *Z,Z,Z,s,s,a* configuration, and c) the green-absorbing photoproduct showing the chromophore in *Z,Z,E,s,s,a* configuration). Note the changed relative orientation of the PCB molecule in b) and c) with respect to helix α4 and beta sheets β5, β6. Electron densities are contoured at 1 σ.

**Figure 8:**
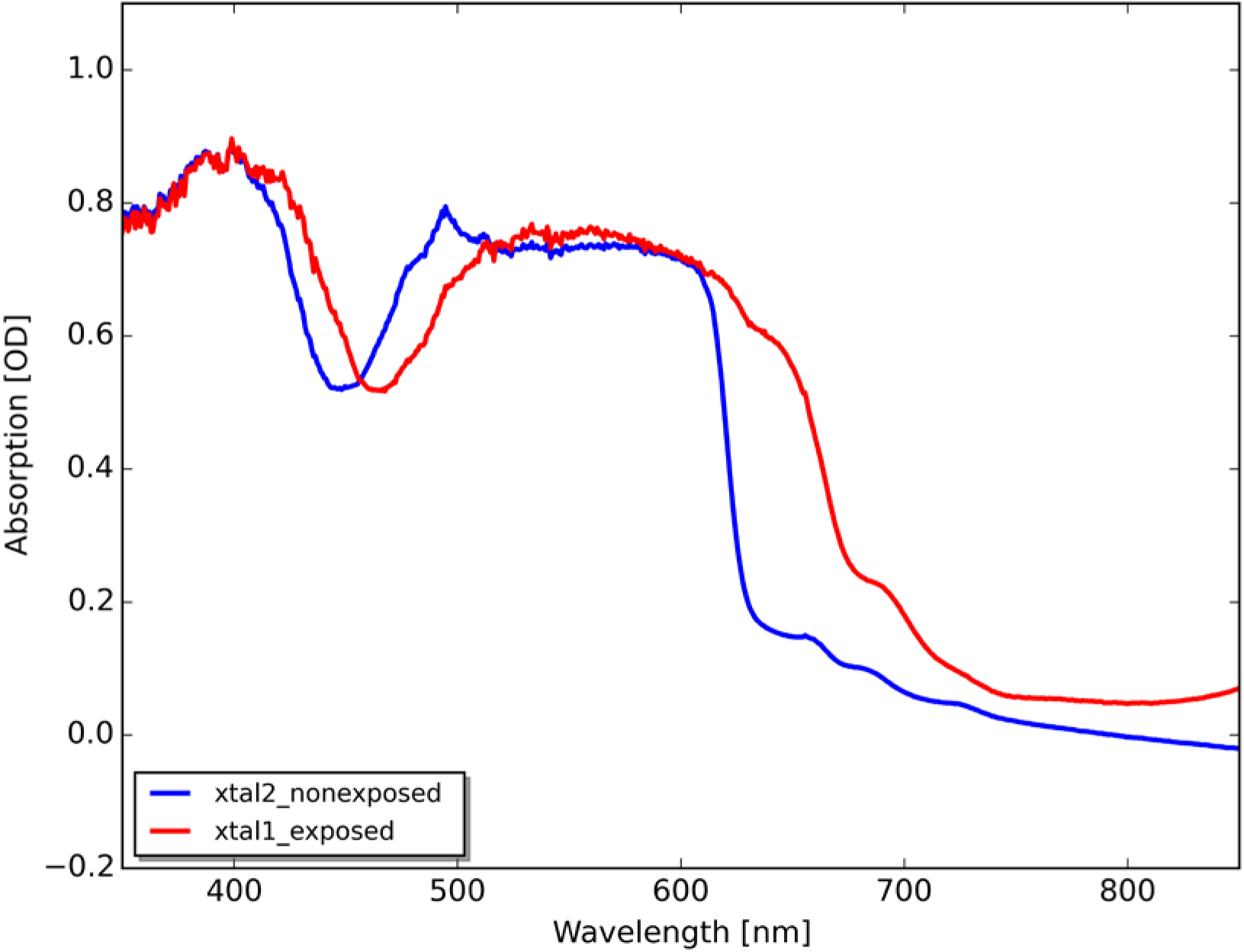
Absorption of Slr1393g3-Pg crystals before (blue trace) and after (red trace) X-ray exposure. Note the growth of absorption around 650 nm after X-ray exposure documenting partial formation of the chromophore in the Pr-state configuration.

Crystals of the intermediate state of Slr1393g3 diffracted to 2.1 Å and contain one monomer per asymmetric unit (ASU). The entire sequence of Slr1393g3 could be modeled into the electron density (residues 441-596) with three C-terminal residues of the expression tag (Glu597, Leu598, Glu599). The R-values after refinement were R_work_ 17.0 % and R_free_ 22.2 %, respectively. The protein fold of the intermediate is nearly identical to the Pg-state (rmsd 0.45 Å over 157 Cα atoms), likewise with the backbone from residue 486 to 499 shifted by *ca*. 8.5 Å towards helix α4 and the additional α-helix from aa 486-492 (α2’). The PCB chromophore is also further shifted into the protein cleft and positioned closer to helix α4 compared to the Pr-state, but shows a somewhat smaller lateral rotation. However, as seen from the clear electron density around the chromophore, the PCB molecule adopts the 15-16 *cis*-conformation, i.e. a *Z,Z,Z,s,s,a* configuration (Figure 7 b).

Despite the different C15-C16 isomeric state, the hydrogen bonding pattern between the PCB chromophore and the Slr1393g3-intermediate protein is similar to the photoproduct Pg-state with few variations: ring A is hydrogen bonded to Thr499 and Asp498, ring B to Arg508, Asp498, one water, one sodium ion and a CAPSO molecule (from the precipitant), ring C interacts with His529, Asp498, one Tris molecule and one sodium ion, whereas ring D interacts only to 4 water molecules (Table 2; Figure S3 c). Like in the Pg-structure, the connector of β2 and α3 is still in the photoproduct conformation and thereby different than in the Pr-state. Thus, Trp496 remains oriented outwards and in close proximity to Cys528 (more than 10 Å apart from the D ring) being out of reach for any π-π stacking. The positionally conserved tryptophan in the ortholog protein AnPixJg2 (Trp289, note that this residue has been assigned as Trp90 in the structural assignment by Narikawa *et al*., 2013) had been suggested to change position upon red-to-green conversion of that protein based on Molecular Dynamics Simulations (Velazquez Escobar et al., 2013). These authors proposed a sideways movement of Trp289 resulting in an inflow of water molecules into the binding pocket of the photoproduct state as a potential explanation for the hypsochromic shift of the absorption peak.

Also in this Slr1393g3-intermediate form the chromophore shows remarkably distortions along its bridging single bonds yielding a conformation that is clearly different to either those of the Pr- or the Pg-state: C4=C5-C6-NB: -62° (pdb labels: C4C=CHD-CID-ND) ; NB-C9-C10=C11: 11° (pdb labels: ND-C4D-CHA=CIA); NC=C14-C15=C16: -36° (pdb labels: NA=C4A-CHB=C1B).

### Correlation with a mutagenesis study by Xu et al., 2014

Our structural data from Slr1393g3, especially from the intermediate and photoproduct state, combined with the parental state structure allow precise identification of conformational changes upon photoisomerization in one and the same phytochrome-related protein with high resolution (drawbacks to the presented structures for TePixJg have already been outlined in the introduction). In addition, the structure now identifies effects formerly reported for site directed mutations (Xu et al., 2014). The mutants described in this manuscript had been classified in three groups, according to their absorbance properties: (i) showing absorbance peaks comparable to the wildtype protein, (ii) exhibiting significantly shifted absorbance maxima, or (iii) presenting a complete loss of *in vivo* chromophore binding ability (Figure 9).

**Figure 9:**
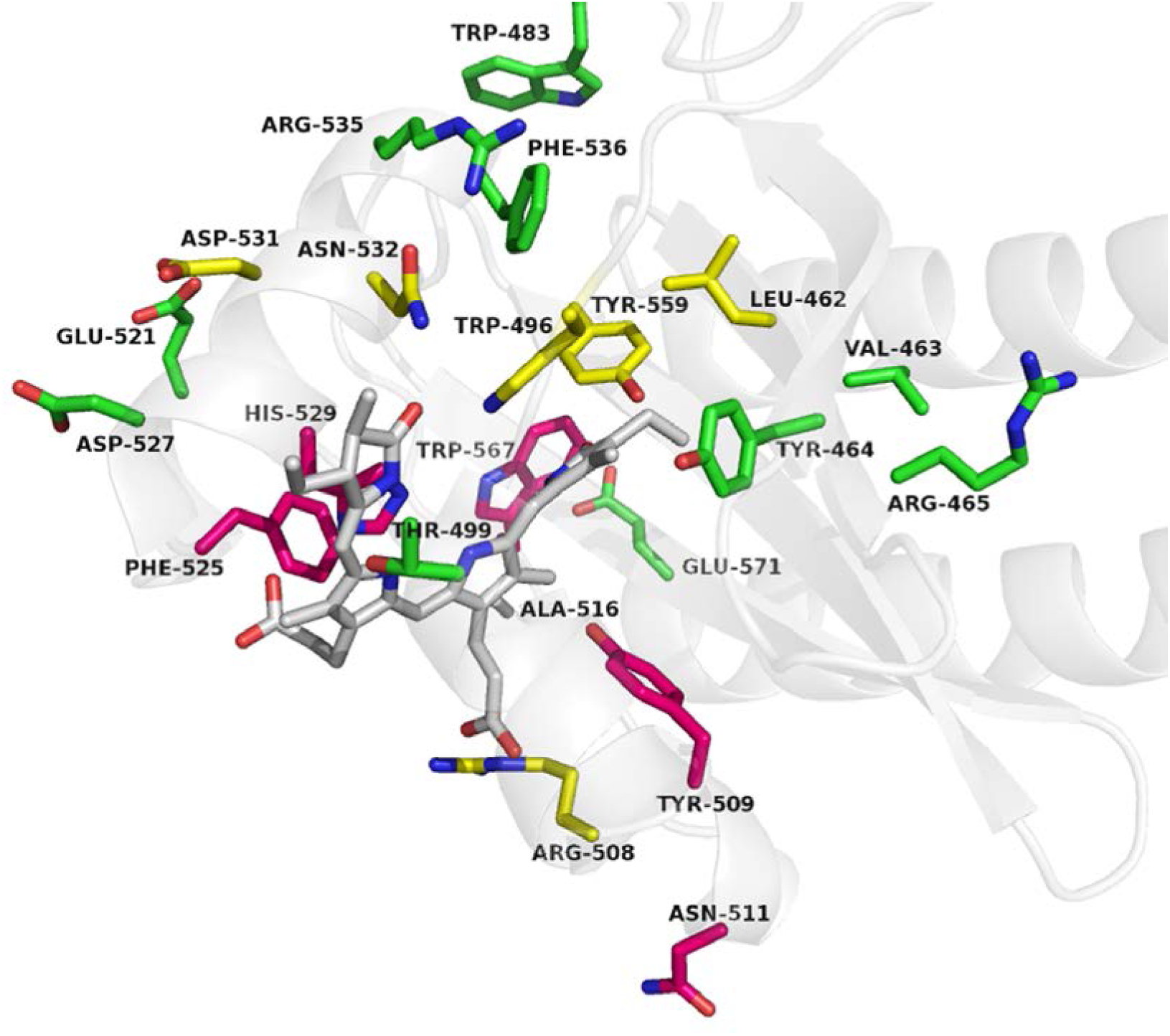
Mutants generated in Xu et al., Innocent’ mutations are shown in green, mutations causing changes in absorption properties are given in yellow, and mutations lacking chromophore assembly are shown in pink. PCB chromophore is depicted in gray.

The first group comprises mutants V463Q, Y464H, R465E, W483M, T499V, A516C/K, E521Y, D527H/I, R535N, F536M and E571A. Except for T499 none of these residues is directly involved in chromophore interactions, neither in the Pr-, nor in the Pg- or in the intermediate state. Almost all sidechains maintain their positions and orientations for each structure resolved, only R535 in Pr-state is pointing in direction to ring D of PCB, but still at a distance of *ca*. 12 Å (in Pg- and intermediate state R535 is oriented outwards on the protein surface). T499 is in hydrogen bonding distance to the chromophore (A-ring, ~3 Å) only in the Pg- and the intermediate state and represents a rather weak interaction, probably insufficient for a strong impact on the spectroscopic properties.

Members of the second group, mutants with significantly shifted spectral properties, are mutants L462S (649, 533 nm), W496I (623, 539 nm), R508N (649, 580 nm), D531T (647, 553 nm), N532Y (651, 546 nm) and Y559H (636, 553 nm); the absorption maxima for the red- and the green absorbing forms are given in brackets (wildtype protein: 648, 538 nm). In the parental and intermediate state of Slr1393g3 the sidechain of Leu462 is positioned around 4.7 Å apart from PCB ring D. In the photoproduct state (Slr1393g3-Pg) this sidechain is in nearly identical position and orientation, but much closer to the D-ring of PCB due to the movement/rotation of the chromophore (~3.8 Å). Therefore, Leu462 might stabilize the chromophore through hydrophobic interactions, which are removed upon mutation to Ser. Trp496, although far apart from the chromophore in the intermediate and Pg-state, is very important for the stabilization of the PCB, in particular for the D-ring via π-π stacking in the Pr-state, as has already been reported for other CBCRs (Velazquez Escobar et al., 2013), and the effect observed upon mutation to Ile especially in the Pr-state now becomes clear. Likewise is the situation for Asn532. Although the position of its sidechain is only slighly different in the Pr- and the intermediate/Pg-state, Asn532 is involved in hydrogen bonding to the A-ring of PCB in the parental state only, since here the chromophore is located more inside the protein cleft and therefore closer to Asn532. Asp531 is not in hydrogen bonding distance to the chromophore in any of the solved structures, but in short distance to the chromophore binding Cys528 in the parental state, i.e. the backbone of Asp531 is involved in a hydrogen bond (~2.8 Å) with Asp527. Mutation from Asp to Thr at position 531 might affect this interaction, resulting in a different orientation of Cys528.

As mentioned before, Arg508 is an important binding partner for the PCB chromophore in all three conformations solved. It interacts either with the propionate group from ring C (parental state) or from ring B in the photoproduct and intermediate state, therefore a mutation to Asn obviously impairs these interactions. The last amino acid in this group of mutants is Tyr559 being one of the D-ring binding residues in the Pr-state. Accordingly, substitution by His extends the distance of the His sidechain to the D-ring (~3.9 Å from 2.6 Å for Tyr) and thus reduces the stabilizing forces.

The third group comprises all mutated variants that have lost the ability of *in vivo* chromophore binding (Y509P, N511K, F525A, H529Y, and W567E). Some of these mutants can be furnished with the chromophore *in vitro*, others have lost also this potential. Tyr509 is not directly involved in chromophore binding, but is in proximity to ring B in the parental state. Tyr509 is anchored in helix α3’, but the mutation T509P, introducing the rigid proline, likely destroys this helical arrangement. Phe525 is located in the loop connecting α4’ and helix α4. Also this amino acid is also not directly involved in chromophore interaction, but Phe525 might act as a sensor for the PCB molecule due to its proximity to the propionate group from ring B in the Pr-state. An effect is also reported upon mutating Trp567 into Glu. Trp567 interacts with the backbone of Val517 and Phe541 and might therefore play a role in stabilizing this part of the β-sheet. Mutation of Asn511 into Lys presents an interesting case. This amino acid is located on the surface of the GAF3 domain with the sidechain pointing outwards. For Slr1393g3-Pr-state we determined a dimeric assembly in solution, and in the crystal Asn511 is forming an H-bond to the backbone oxygen of Lys487, wherefore the Asn-to-Lys mutation at this position might change the dimer interface due to the much longer sidechain of Lys (Figure S2 a). The last amino acid in this group, His529, is one of the most important binding residues for the PCB chromophore in all states. In the parental state it interacts with the propionate group from ring B, but in the Pg- and intermediate state with the propionate group from ring C. Replacement of this sidechain by Tyr (H529Y) most likely causes two different effects both leading to the loss of chromophore binding capacity: the interaction with either of the propionate side chains is eliminated and the longer sidechain induces sterical hindrances within the binding pocket, reducing the available space for the PCB molecule. This is in good agreement with the fact that the mutant H529Y cannot be furnished with the chromophore, neither *in vivo* nor *in vitro*.

### Factors affecting the absorption properties in phytochromes and related proteins

In addition to the direct comparison of two states of Slr1393g3, the structural determination of a photoisomerization intermediate with the protein still in the photoproduct state and the chromophore already in 15-*Z* conformation of the parental state clearly adds to our knowledge of chromophore-protein interaction in phytochromes. However, our findings also challenge proposed mechanisms that regulate the absorbance of the various states. We find a color change of intermediate state crystals (at positions where the crystals were exposed to the X-ray beam) concomitant with the chromophore in 15-*Z* configuration (λ_max_ around 650 nm, Figure 8), despite the presence (parental state) or absence (photoproduct) of the π-π stacking with Trp496, and irrespective of an altered protein surrounding and interactions with amino acids and water molecules. Apparently, it is not necessary that the rearrangement of the binding site offers a more polar environment to the chromophore causing a color change (as was proposed on the basis of quantum mechanical calculations (Velazquez Escobar et al., 2013)), but apparently the absorption properties are intrinsic to the conformation, respectively the distortion along bridging single bonds, of the chromophore itself (with its particular interactions to amino acid side chains in its close proximity). Accordingly, both 15-*Z* and 15-*E* isomers of Slr1393g3 show A- and D-rings in strong, but different distortion angles along the bridging single bonds. This is highlighted by an overlay of the chromophore conformation in all three structurally resolved states of Slr1393g3 described here, the Pr-, the Pg-, and the intermediate state (Figure 10, with either, Figure 10 a, having the A-ring in front, or, Figure 10 b, having the D-ring in front).

**Figure 10:**
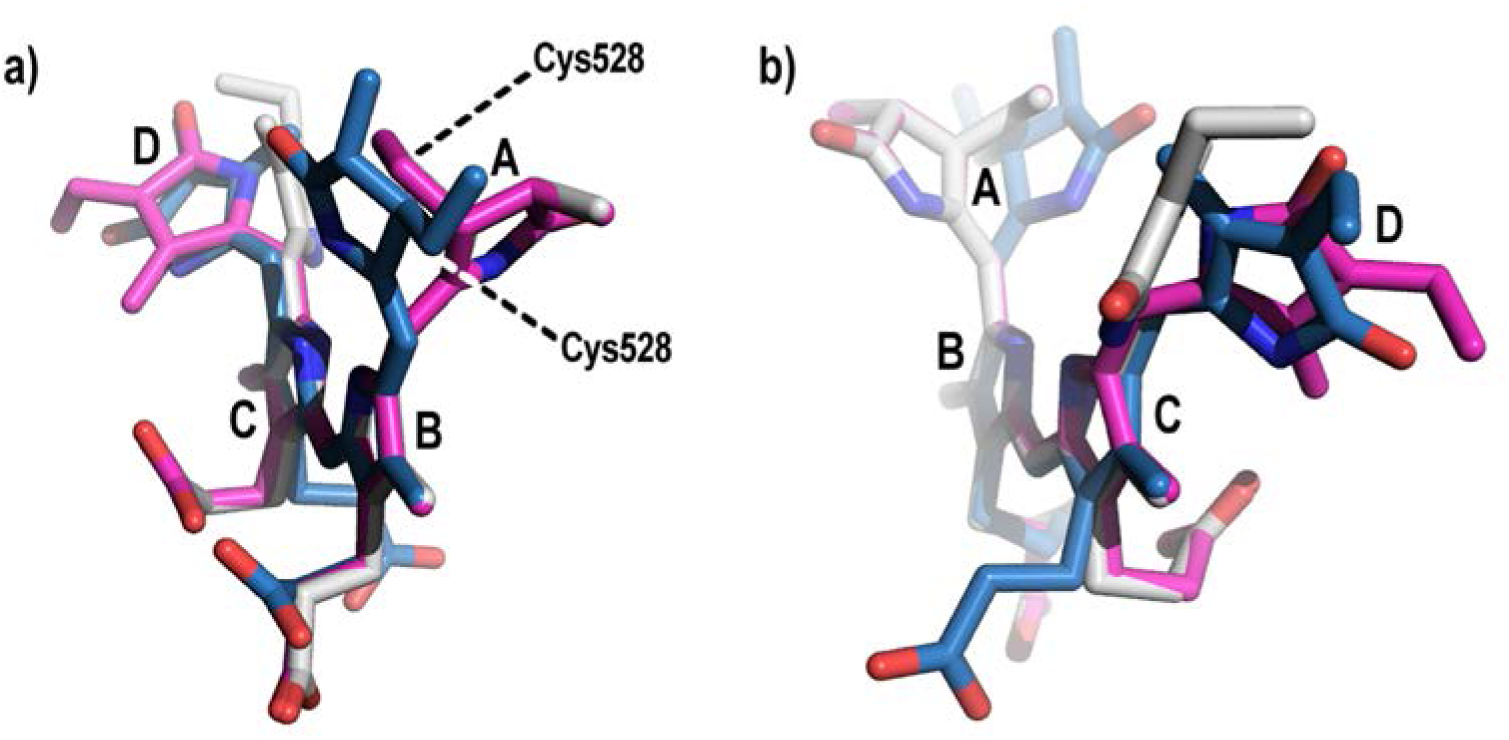
Superposition of the PCB chromophores of Slr1393g3 in Pr-state (blue, 5DFY), intermediate state (gray, 5M85) and Pg-state (magenta, 5M82). Ring B and C were used as reference points. a) shows the chromophore conformation along rings B-C, b) along rings C-B. Remarkably are the two different directions of distortion of ring A with respect to the plane spanned by rings B-C in Pr-state versus Pr-/intermediate state.

It is intriguing to compare chromophore conformations in various crystallized phytochromes in order to shed light onto structural arrangements that guide the absorption position of the phytochrome(- related) proteins, however, many structures provide a resolution too low for precise chromophore conformation determination, and as such, dihedral angles sometimes reside more on model building (for a list of phytochromes and related proteins with reasonable resolution see Table S1). In general, the dihedral angle between rings B and C is smallest, reflecting the high degree of conjugation in these two rings for ‘relaxed’ chromophores in their parental state; it resides between -0.8° and *ca.* 10° for ‘canonical’ phytochromes (Cph1 from *Synechocystis* (2VEA), *A. thaliana* PhyB (4OUR), and the bacteriophytochromes from *D. radiodurans* (4Q0H) and *S. aurantiaca* (4RPW) with 2.3°, 0.5°, 3.2°, and 2.9°), but near planarity is also found for AnPixJg2 from *Anabaena* (2.2°, 3W2Z) and Slr1393g3 from *Synechocystis* (-0.8°, this work). Common to these proteins is also the large negative dihedral angle between rings C and D (negative values arise from the *anti* configuration of the 14-15 single bond), all found in a range of *ca.* 150° and 160°. For these proteins, a larger variation (between 0° and ca. 20°) is found for the dihedral angle between rings A and B. Stronger deviations from these ranges of dihedral angles are seen for phytochromes with unusual properties, e.g., bathy-phytochrome from *P. aeruginosa* (3NHQ) with an unusually large B-C ring angle (17°) or the cyanobacteriochrome TePixJg (4FOF, 3VV4), switching between a blue- and green-absorbing state. For this protein strongly tilted rings are found for both states: -96.4°, 11.3°, and -134.6° (A-B, B-C, and C-D) for the green-absorbing state and -134.4°, -69.2°, and -162.9° for the blue absorbing state. Similarly strong deviations from the range of phytochromes with ,relaxed’ conformation are also found for the Slr1393g3 15-*E* state (green absorbing) and the intermediate form (both this work): - 61.1°, 9.2°, and -110° for Pg, and -35.7°, 7.0°, and -61.8° for the intermediate. Clearly, the experimental basis to extract a mechanism by which the absorption band position is regulated is still too small, however, one may propose that strong distortions along the A-B bridge (still with the C4=C5 double bond in *Z*) point to strongly shifted absorption maxima.

## Conclusion

The crystallization of cyanobacterial bilin-binding GAF3 domain from Slr1393 in both parental and photoproduct state with high resolution allows a direct comparison of conformational changes induced by light in one and the same protein. This situation is advantageous over many other comparative studies that employ two phytochrome-related proteins, for which either the 15-*Z* (canonical phytochromes) or the 15-*E* isomer (bathy-phytochromes) is the parental state. Clearly, such approaches suffer from comparing proteins with different sequences and potentially different arrangement of secondary structural elements. Despite the overall well-preserved tertiary structure of both Slr1393g3 states, significant conformational changes are precisely identified: whereas the parental state shows a long unstructured loop extending over the chromophore, part of this loop is formed into a small α-helix, thus adding structural stabilization to the photoproduct’s chromophore conformation. Importantly, a tryptophan residue (Trp496) as part of this loop, being in π-π stacking arrangement with ring D in the parental state, is moved away from the isomerized chromophore by ca. 14 Å, opening the binding pocket to the external medium. The chromophore isomerization causes also a rotation of the PCB molecule around its central orthogonal axis such that Arg508 (interacting amino acid to the propionate group of ring C in the parental state) now hand-shakes with the propionate group of ring B, concomitant with a reduction of the distance between chromophore and helix α4. The structures of the intermediate and the Pg-state of Slr1393g3 in combination with absorption spectra of both crystals indicate that the absorption properties are intrinsic to the conformation of the chromophore itself, respectively the distortion along bridging single bonds, whereas the protein environment has an only marginal effect.

## Materials and Methods

### Protein preparation

GAF3 domain of Slr1393 from the cyanobacterium *Synechocystis* PCC6803 has been prepared and purified as recently described, following a two-plasmid expression protocol that yields protein and chromophore *in vivo* allowing formation of the holoprotein during biosynthesis (Chen et al., 2012; Xu et al., 2014). In brief, Slr1393g3-transformed *E. coli* cells were harvested after IPTG induction by centrifugation and lysed in liquid nitrogen by treatment with an Ultraturrax. The soluble fraction was subjected to IMAC for isolation of the His-tagged CBCR-GAF domain. Pooled protein fractions were further purified via anion exchange chromatography. As a proof of photochemical homogeneity, samples of pure CBCR-GAF3 were irradiated with appropriate light using LEDs yielding both parental and photoproduct state to nearly 100 %. Alternatively, solely the protein moiety was expressed and purified by IMAC. After this first purification step, PCB was added in small portions until no further increase at 650 nm was observed. Second step purification by anion exchange chromatography was as for the *in vivo* prepared protein.

### Crystallization conditions and Structure Determination

Slr1393g3*-Pr-state, in vivo:* Initial crystals were optimized using sitting drop vapor diffusion method with drops containing of 1 μl of protein solution (10 mg/ml in 50 mM Tris-HCl pH 7.5, 100 mM NaCl) mixed with 1 μl reservoir solution (0.02 M Tris-HCl pH 7.0, 0.1 M sodium chloride, 7.7 % (w/v) PEG 4000), equilibrated over 300 μl reservoir at 12 °C. Crystallization experiments were set up under dim green-light and crystallization plates were covered with aluminum foil. Blue rhomboid-shaped crystals grew to a final size of 100 × 100 × 70 μm^3^ within 8 weeks. Crystal-containing drops were overlaid with mineral oil for cryo-protection before harvesting the crystals. Crystals were shock-frozen with liquid nitrogen.

Slr1393g3*-Pr-state, in vitro*: Crystallization trials and crystal harvesting was performed as described above but under different crystallization conditions and in hanging drops. The optimized precipitant contained 0.02 M sodium citrate pH 5.6, 0.1 M sodium chloride and 11 % (w/v) PEG 3350. Blue rhomboid-shaped crystals grew to a final size of 120 × 120 × 100 μm^3^ within 8 weeks. Crystal-containing drops were overlaid with mineral oil for cryo-protection before harvesting the crystals. Again, crystals were shock-frozen in liquid nitrogen.

Slr1393g3*-Pg-state* and *-intermediate-state*: Prior to crystallization experiments the protein solution was irradiated with appropriate light using red LEDs to generate the photoproduct state. All crystallization experiments were set up under dim red-light. Initial crystals were optimized using sitting drop vapor diffusion method with drops containing of 1 μl of protein solution (8-10 mg/ml in 50 mM Tris-HCl pH 7.2, 100 mM NaCl) mixed with 1 μl reservoir solution (0.1 M CAPSO pH 9.5, 0.1 M sodium chloride, 100-175 mM lithium sulphate, 12-14 % (w/v) PEG 4000), equilibrated over 300 μl reservoir at 4 °C. The crystallization plates were kept under red light. Pink rhomboid-shaped crystals grew to a final size of 90 × 90 × 60 μm^3^ within 2 weeks. Crystal-containing drops were overlaid with mineral oil for cryo-protection before harvesting the crystals. Crystals were shock-frozen with liquid nitrogen. All steps were performed under dim red-light.

### Data collection

Slr1393g3*-Pr-state* (*in vivo*): Data were collected at 100 K from a single crystal at ID29, ESRF, Grenoble, France. Data were processed by XDS and XSCALE (Kabsch, 2010b; Kabsch, 2010a). The structure was solved by molecular replacement (MR) using the autorickshaw pipeline (Panjikar et al., 2005) providing the protein sequence and processed data as input files. The structural model was further built and refined using iterative cycles of manual building in coot (Emsley et al., 2010) and refinement cycles using REFMAC5 (Murshudov et al., 2011) from the ccp4 suite (1994)(Collaborative Computational Project, 1994). The structure was deposited in the protein data bank (PDB) under the accession code 5DFX.

Slr1393g3*-Pr-state* (*in vitro*): Data were collected at 100 K from a single crystal at ID23-1, ESRF, Grenoble, France. Data were processed by XDS (Kabsch, 2010b; Kabsch, 2010a). The structure was solved by molecular replacement (MR) using 5DFX as search model. The structural model was further built and refined using iterative cycles of manual building in coot (Emsley et al., 2010) and refinement cycles using REFMAC5 (Murshudov et al., 2011) from the ccp4 suite (Collaborative Computational Project, 1994). The structure was deposited in the protein data bank (PDB) under the accession code 5DFY.

Slr1393g3*-intermediate-state*: Data were collected at 100 K from a single crystal at P13, DESY, EMBL, Hamburg, Germany (Cianci et al., 2017). During data collection the crystal area exposed to x-rays immediately turned from pink to dark blue indicating a conformational change in the chromophore arrangement. Data were processed by XDS (Kabsch, 2010a, b). The structure was solved by molecular replacement (MR) using 5DFX as search model. The structural model was further built and refined using iterative cycles of manual building in coot (Emsley *et al.*, 2010) and refinement cycles using REFMAC5 (Murshudov *et al.*, 2011) from the ccp4 suite (Collaborative Computational Project, 1994). The structure was deposited in the protein data bank (PDB) under the accession code 5M85. Slr1393g3*-Pg-state*: Data were collected at 100 K from a single crystal at P13, DESY, EMBL, Hamburg, Germany (Cianci *et al.*, 2017). As experienced from the pink crystals used for determination of the intermediate state we knew the crystal area exposed to x-rays was highly susceptible to turn from pink to blue, we here used a helical data collection strategy with minimum exposure time and minimum total oscillation to reduce the effect of the x-rays to the chromophore arrangement. Data were processed by XDS (Kabsch, 2010a; Kabsch, 2010b). The structure was solved by molecular replacement (MR) using 5DFX as search model. The structural model was further built and refined using iterative cycles of manual building in coot (Emsley et al., 2010) and refinement cycles using REFMAC5 (Murshudov et al., 2011) from the ccp4 suite (Collaborative Computational Project, 1994). The structure was deposited in the protein data bank (PDB) under the accession code 5M82.

Detailed information about all data collection and refinement statistics are given in Table 1.

All images of the models were prepared using PyMOL (The PyMOL molecular graphics system). 2D ligand plots of interactions were generated using the webservice „PDBsum Generate“ (http://www.ebi.ac.uk/thornton-srv/databases/pdbsum/Generate.html).

### Accession Numbers

The crystal structure of Slr1393g3 is deposited at PDB (wwpdb) under 5DFY and 5DFX (*in vitro* and *in vivo* assembled) in the dark (red-absorbing) state, under 5M82 in the light state (green-absorbing), and under 5M85 in the intermediate state.

## Supplemental Material

Table S1, Figures S1 – S4.

## Acknowledgements

The expert technical help from Stefanie Kobus, X-ray Facility and Crystal Farm, Heinrich-Heine-Universität, is greatly acknowledged. We thank Sander Smits (Institute of Biochemistry, Heinrich-Heine-Universität Düsseldorf) for helpful discussion and advice during development of data collection strategies. We acknowledge the European Synchrotron Radiation Facility for provision of synchrotron radiation facilities and we wish to thank Antoine Royant and Ulrich Zander for assistance in using beamline ID23-1 and ID29. We also thank the Isabel Bento and Guillaume Pompidor at P13, DESY (EMBL, Hamburg, Germany) for kind support during data collection. KHZ is supported by grants 21472055 and 31270893 from the National Natural Science Foundation of China. WG gratefully acknowledges the generous financial support from the Max-Planck-Society.

## Declaration

The authors declare that there is no financial or non-financial competing interest.

## Supplemental Material

**Table S1:** Crystal structures of phytochromes and CBCR proteins (from PDB database) with resolution <3.5 Å.

| Name<sup>1</sup> (state) | pdb code | res. (Å) | A-B (°) | B-C (°) | C-D (°) | Notes<sup>2</sup> |
| --- | --- | --- | --- | --- | --- | --- |
| Cph1 (Pr) | 2VEA | 2.21 | 13 | 2.3 | -152 |  |
| *At*-phyB (Pr) | 4OUR | 3.4 | 0 | 0.5 | -145 | low res. |
| Cph2 (Pr) | 4BWI | 2.6 | 31.44 | 7.2 | -144 | res. limit |
| *Te*-PixJ (Pg)<sup>4</sup> | 3VV4 | 2.0 | -96.4 | 11.3 | -134.6 |  |
| *Te*-PixJ (Pb)<sup>4,5</sup> | 4FOF | 2.42 | -134.4 | -69.2 | -162.9 |  |
| *An*-PixJ (Pr) | 3W2Z | 1.8 | 10.6 | 2.2 | -148 |  |
| *Pa*-bathy-phy (Pfr) | 3NHQ | 2.55 | 10.7 | 17.2 | -149 | res. limit |
| *Dr*-bacterio-phy (Pr) | 5K5B | 1.35 | 11.3 | 3.0 | -156.8 |  |
| *Dr*-bacterio-phy (Pr) | 4Q0H | 1.16 | 1.64 | 3.24 | -150.2 |  |
| *Dr*-bacterio-phy (Pfr) | 5C5K | 3.31 | 15.14 | 0 | -145.7 | low res. |
| *Sa*-bacterio-phy (Pr) | 4RPW | 2.73 | -2.34 | 2.89 | -156.7 | res. limit |
| *Xcc*-bacterio-phy (Pr) | 5AKP | 3.25 | 12.2 | 6.34 | -159 | low res. |
| Slr1393g3 (Pr) | 5DFY | 1.6 | 21 | -4 | -155.6 | this work |
| Slr1393g3 (Pr) | 5DFX | 1.8 | 23.8 | -0.8 | -150.6 | this work |
| Slr1393g3 (Pg) | 5M82 | 1.9 | -61.1 | 9.2 | -110 | this work |
| Slr1393g3 (Pi)<sup>3</sup> | 5M85 | 2.1 | -35.7 | 7.0 | -61.8 | this work |

<sup>1</sup>, this table does not contain NMR-based structures
<sup>2</sup>, definitions were set as of: resolution > 2.5 Å: resolution limit; resolution > 3 Å: low resolution
<sup>3</sup>, isomerization intermediate
<sup>4</sup>, TePixJ autocatalytically converts its PCB chromophore into phycoviolobin (PVB)
<sup>5</sup>, The blue-absorbing, short wavelength absorption is accomplished by covalent attachment of cysteine residue to carbon 10 of PVB thus restricting the $\pi$ -electron conjugation to rings C and D.

**Figure S1:**
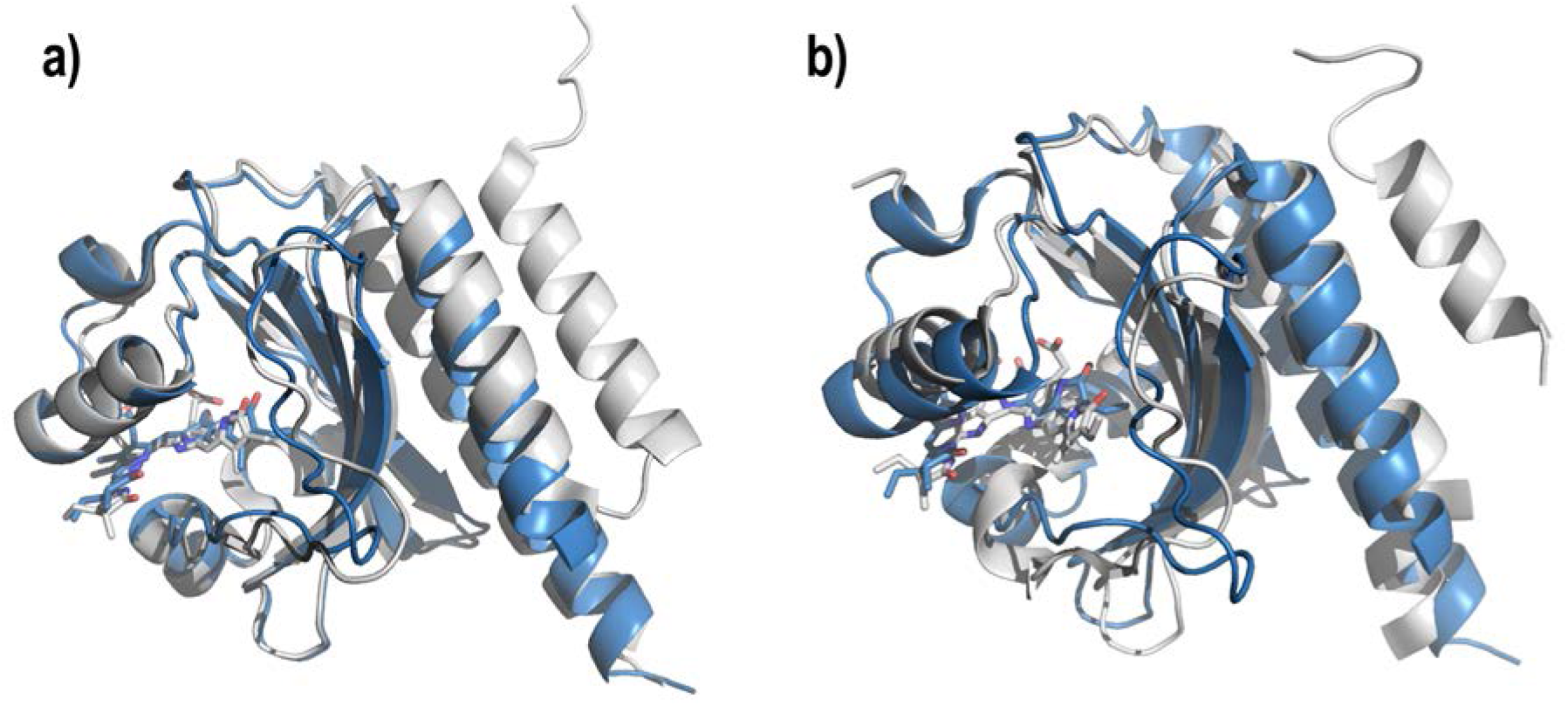
Superposition of the Slr1393g3-Pr structure (5DFY, blue cartoon) and a) AnPixJg2 (3W2Z, gray cartoon) and b) the GAF domain of Cph1 (2VEA, gray cartoon).

**Fig. S2:**
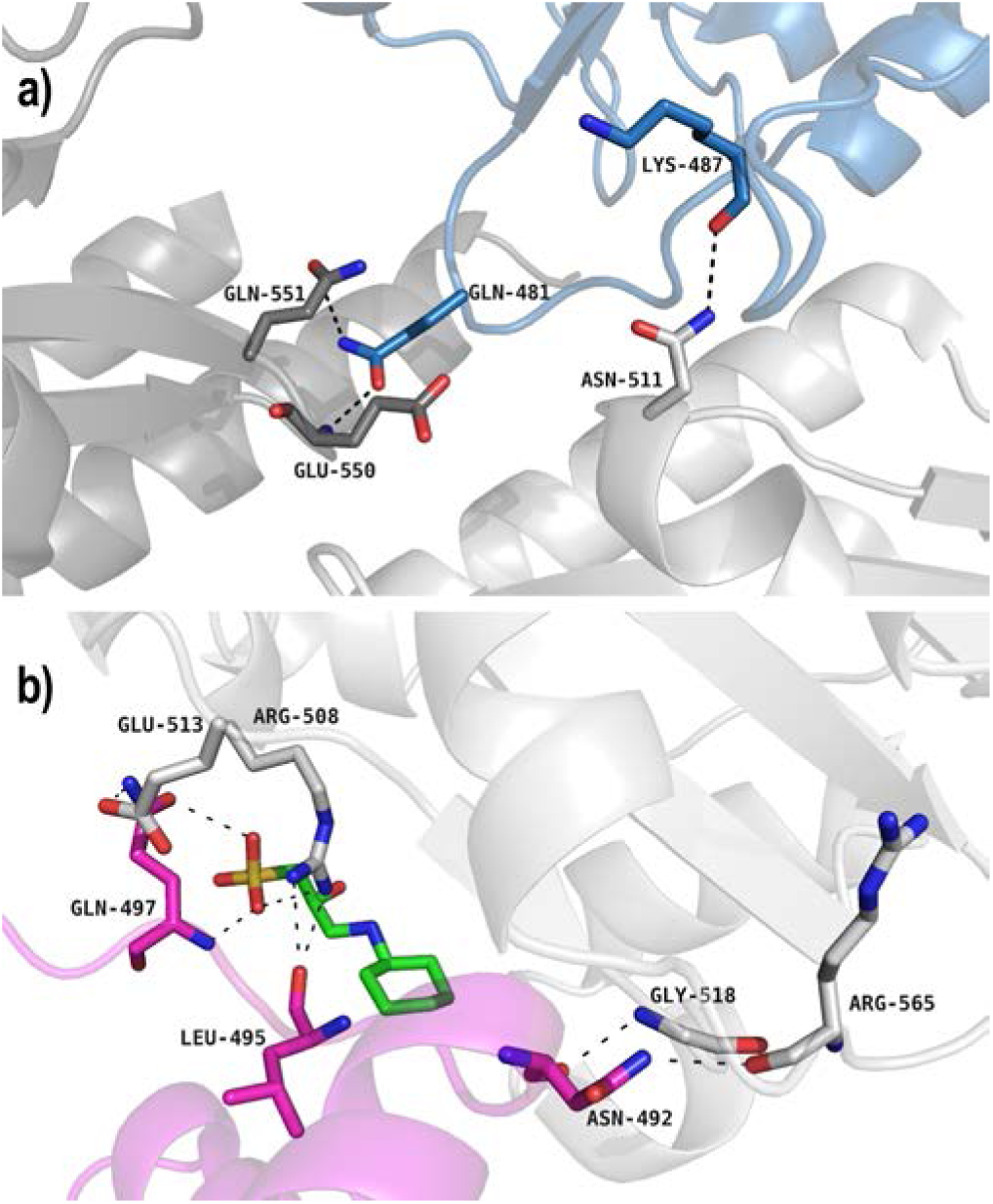
Intermolecular crystal contacts for a) the Pr-state (5DFY) and b) the Pg-state of Slr1393g3 (5M82). The protein backbones are represented by cartoons, the residues involved in interactions are depicted as sticks. In a) three monomers are shown (blue, dark gray, light gray), in b) two monomers (magenta, light gray). In the Pg-state a CAPSO molecule is part of the bridging network (green sticks).

**Figure. S3:**
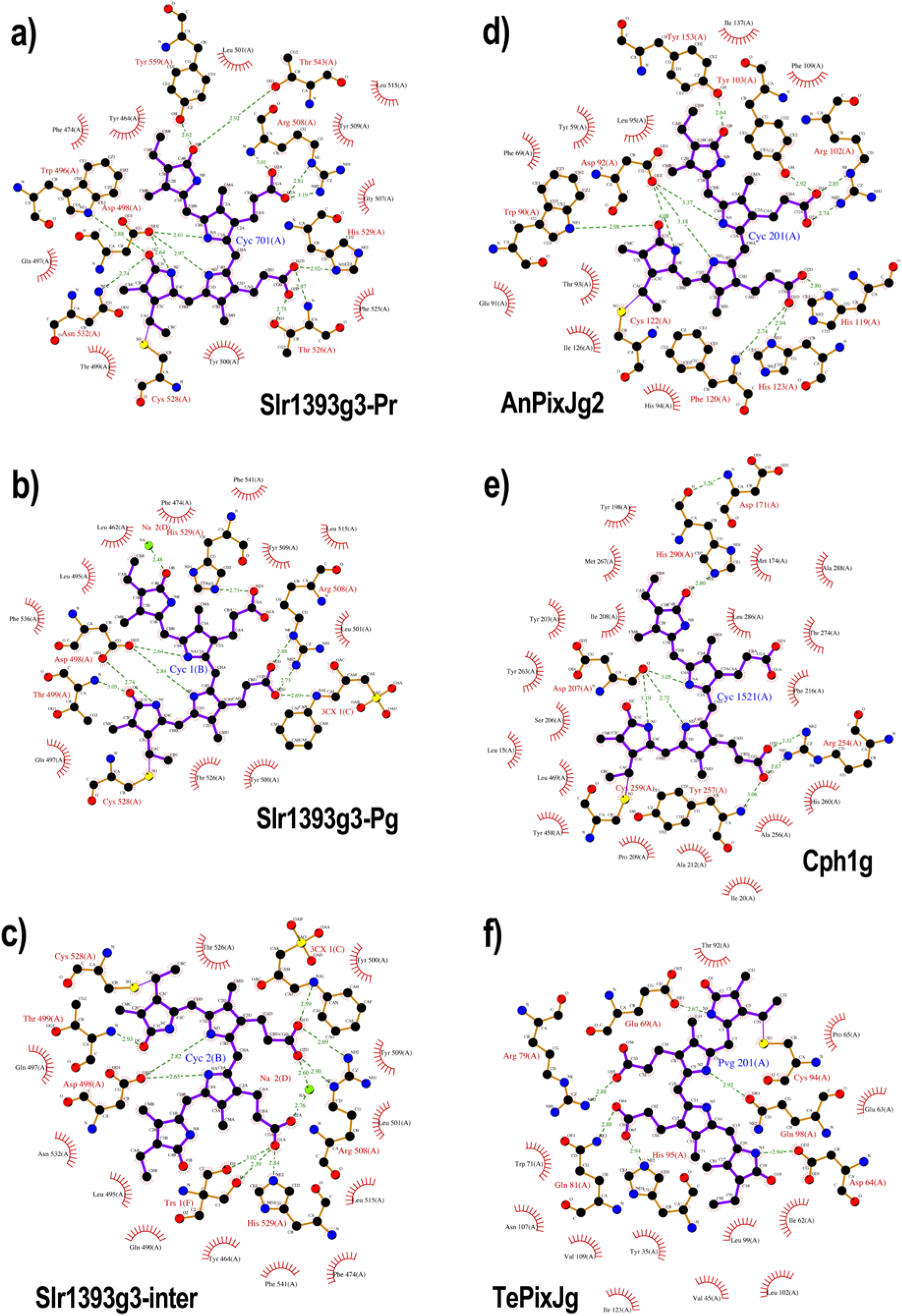
Comparison of chromophore-protein contacts, represented by 2D ligand plots. a), b) and c) are Slr1393g3 structures with a) the red-absorbing parental state (5DFY), b) the green-absorbing photoproduct state (5M82), and c) the photointermediate (5M85); d) AnPixJg2 (3W2Z), e) Cph1 (2VEA) and f) TePixJ (3VV4). The chromophore molecules and interacting residues are represented as ball-and-sticks, distances are given in Å (dotted lines). Residues in close proximity to the chromophore, but not in hydrogen bonding distance are depicted as semi-circles, water molecules are omitted. Note that these plots are purely schematical representations of the interactions and do not reflect absolute chemical configurations.

**Figure S4:**
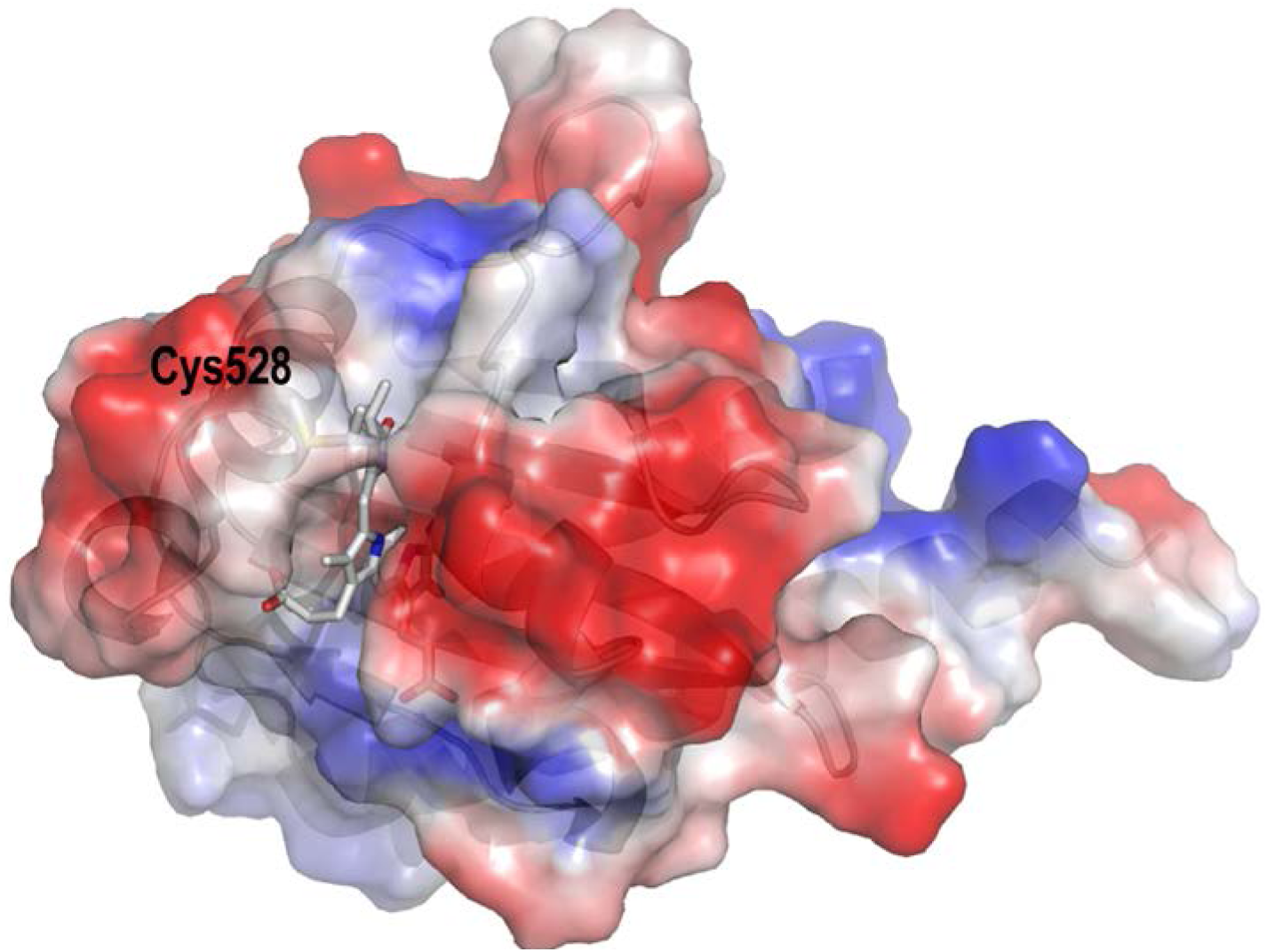
Surface charge of the parental state of Slr1393g3 (5DFY). For clarity the protein is shown as cartoon, the PCB chromophore and the covalent binding to Cys528 are depicted as gray sticks. Red: acidic region, blue: basic region.

